# Global multi-omics analysis of exerkines in progressive treadmill-exercised rodents

**DOI:** 10.1101/2024.12.08.627439

**Authors:** Hash Brown Taha

## Abstract

Exercise is widely recognized for its comprehensive physiological benefits, attributed largely to the secretion of signaling molecules known as exerkines. These molecules, originating from various tissues like muscles, brain, and liver, facilitate inter-organ communication and enhance metabolic health, immune function, and tissue repair. However, the responsiveness of multiple tissues and exerkines to the same exercise regimen remains poorly understood. To address this issue and elucidate patterns of time-dependent, intensity-related and sex-dimorphic tissue and exerkine responsiveness, we leveraged the publicly available Molecular Transducers of Physical Activity Consortium (MoTrPAC) dataset. Male and female Fischer 344 rats aged 6 months underwent a progressive treadmill training protocol designed to emulate human endurance exercise. Blood (cells and plasma) and 18 solid tissues such as adipose, skeletal muscle and brain were collected and multi-omics analyses, including proteomics and transcriptomics were performed on them. We examined the distribution of 26 known and 2 speculative exerkines across 2 biofluids and 18 solid tissues. Our analysis reveals that brown adipose tissue (BAT), the adrenal gland, and white adipose tissue (WAT) are the most responsive to exercise-induced changes. Fractalkine was the most responsive exerkine, followed by prosaposin (speculative), cathepsin B, and FNDC5/irisin, platelet factor 4, Clusterin and SPARC. Additionally, we found distinct patterns in the responsiveness of tissues and exerkines based on the duration and intensity of exercise, with notable differences between male and female rodents. Future research should investigate whether our findings on tissue exerkine responsiveness vary with age and disease status, and determine if these findings can be extrapolated to human populations.

## Introduction

Exercise is known to be a panacea with comprehensive physiological benefits for the whole body. Historically, even with limited understanding of the mechanisms driving various diseases, exercise has been recommended for its broad physiological benefits and its ability to prevent and modify diseases, promoting overall health. The widespread health benefits of exercise are largely attributed to the secretion of signaling molecules from tissues during both acute and chronic exercise. These molecules, known as “exerkines,” have autocrine, paracrine, and endocrine effects that impact multiple organ systems [1–3]. Indeed, recent studies demonstrate that re-infusing exercise-conditioned plasma into non-exercised mice confirms the presence of bioactive exerkines in the bloodstream, offering protection against age-related cognitive decline and enhancing memory [4; 5].

Many exerkines have been identified that originate from a variety of tissues including muscles, brain, adipose tissue, liver and the brain, among others [1; 2; 6]. Exerkines play an important role in facilitating inter-organ communication, mediating the systemic effects of exercise on metabolic health, immune function, and tissue repair. For example, myokines such as interleukin-6 (IL-6) are released by muscles and influence glucose metabolism and inflammation. The brain secretes brain-derived neurotrophic factor (BDNF), which supports neuroplasticity and cognitive function. Adipose tissue produces adiponectin, which improves insulin sensitivity, while the liver releases fibroblast growth factor 21 (FGF21) to regulate energy homeostasis. These molecules are not isolated in their action; rather, they may potentially exert synergistic or additive effects to amplify their influence on whole-body metabolic health [1; 7].

Previous studies have mostly focused on studying a single factor (e.g., BDNF) using specific tissues (e.g., brain). It remains unclear which tissue is most responsive to exercise, as well as which ones act synergistically or additively across different organ systems to mediate the health benefits of exercise. Furthermore, it is unclear whether there are time-dependent, intensity-related or sex-dimorphic variations in exerkine expression and function, which are further complicated by differences in exercise regimens, variations in sampled tissues, and quantification methods. Conducting comprehensive and integrative studies to address these issues is challenging due to the high costs, intensive labor, and significant time requirements involved.

The Molecular Transducers of Physical Activity Consortium (MoTrPAC) is a large-scale, global study using both human and rats aimed at uncovering the molecular mechanisms by which exercise benefits health. Male and female Fischer 344 rats aged 6 months underwent a progressive treadmill training protocol designed to emulate human endurance exercise (**Figure 1A**; reproduced from [6]). Blood (cells and plasma) and 18 solid tissues such as adipose, skeletal muscle and brain were collected and multi-omics analyses, including proteomics and transcriptomics were performed on them (**Figure 1B**; reproduced from [6]) to comprehensively map tissue-specific exercise-induced molecular changes [6].

**Figure 1.**
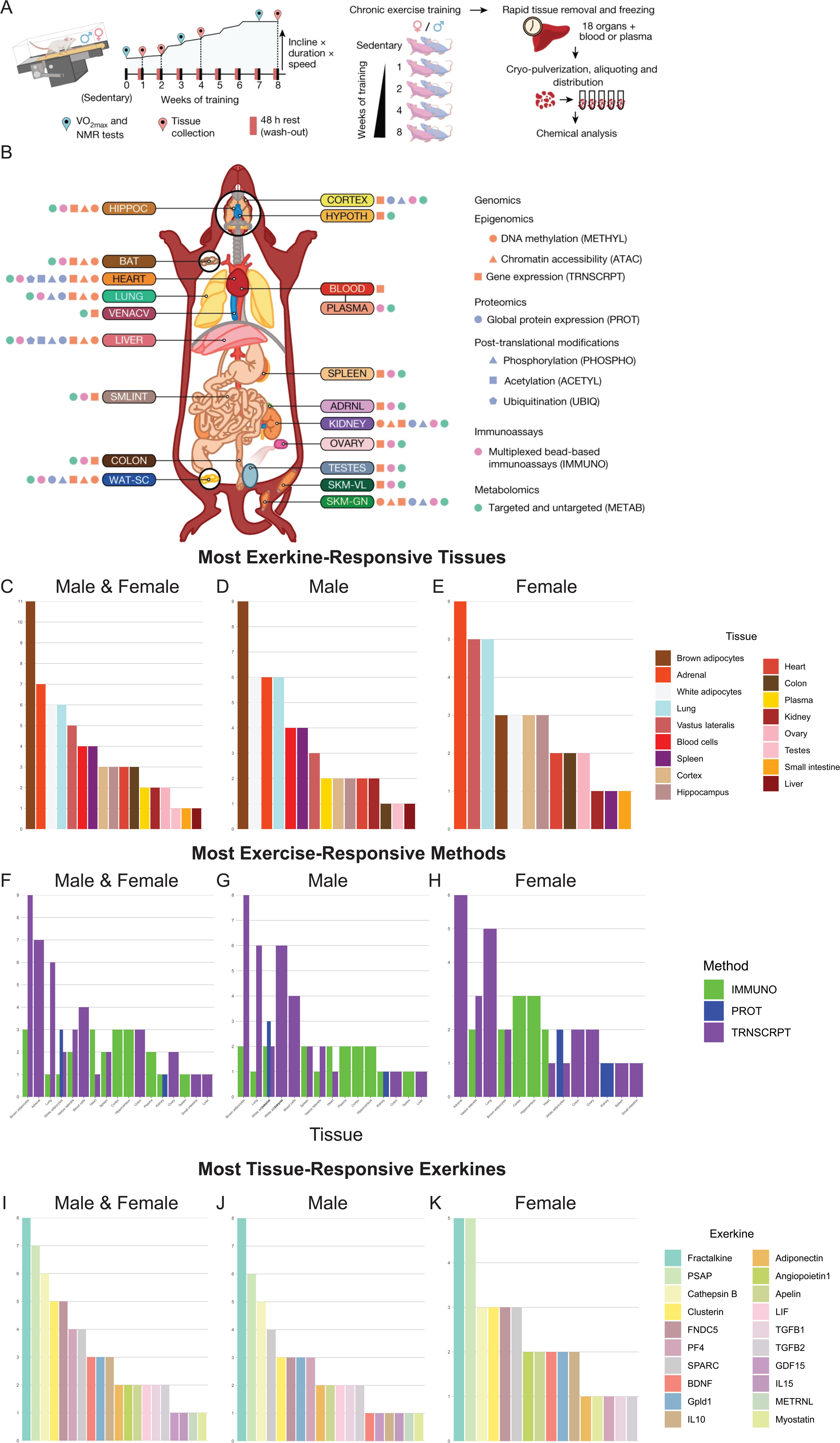
Experimental design and tissue sample processing. Rats were subjected to a progressive treadmill training protocol. Tissues were collected from male and female rats that remained sedentary or completed 1, 2, 4 or 8 weeks of progressive endurance exercise training. For trained animals, samples were collected 48 hours after their last exercise bout (**A**). Summary of molecular datasets included in this study. Up to nine data types (omes) were generated for blood cells, plasma, and 18 solid tissues per animal (**B**). Panels A-B are reproduced from the MoTrPAC [6] study under a Creative Commons Attribution 4.0 International License (CC BY 4.0), which permits unrestricted use, distribution, and reproduction in any medium, provided the original author and source are credited. Distribution of exerkine-responsive tissues (**C-E**), exerkine-responsive methods (**F-H**) and tissue-responsive exerkines (**I-K**).

Herein, we gathered a list of popular exerkines and analyzed their expression using the 6-month MoTrPAC rat dataset [6]. Our objectives were to identify the most exercise-responsive tissues and exerkines, elucidate tissue- and molecular-specific patterns, explore potential inter-organ roles, investigate any time-dependent, intensity-related or sex-dimorphic patterns and make comparisons to the published literature.

## Methods

Data analyzed in this study is publicly available from the MoTrPAC study. Exercise regimens, including progressive treadmill training at 70%–75% of VO max, duration of 1, 2, 4, or 8 weeks, and details on sex-specific (male and female) and age-specific (6-month-old adult and 18- month-old aged rats) cohorts, are comprehensively described in the original study [6].

We analyzed a list of 26 exerkines and two speculative exerkines [1; 4; 5; 8; 9] across blood (cells and plasma) and 18 solid tissues in the MoTrPaC rat 6-month dataset. The exerkines include apelin (APLN), adiponectin (ADIPOQ), meteorin-like (METRNL), FGF21, growth differentiation factor 15 (GDF15), fibronectin type III domain-containing protein 5 (FNDC5)/irisin, cathepsin B (CTSB), IL-6, interleukin 7 (IL-7), interleukin 10 (IL-10), interleukin 13 (IL-13), interleukin 15 (IL-15), musclin/osteocrin (OSTN), myostatin (MSTN), leukemia inhibitory factor (LIF), secreted protein acidic and rich in cysteine (SPARC), syndecan 4 (SDC4), transforming growth factor beta 1 (TGFβ1), transforming growth factor beta 2 (TGFβ2), angiopoietin 1 (ANGPT1), fractalkine (CX3CL1), oxytocin (OXT), BDNF, platelet factor 4 (PF4), glycosylphosphatidylinositol specific phospholipase D1 (GPLD1) and Clusterin (Clu). We also included the speculative exerkines prosaposin (PSAP) and PSAP like 1 (PSAPL1) given that it is gaining attention as a new myokine and adipokine [10; 11]. The solid tissues include the brain (cortex, hippocampus, hypothalamus), skeletal muscle (gastrocnemius and vastus lateralis/quadriceps), heart, kidney, adrenal gland, colon, spleen, testes, ovaries, lungs, small intestine, liver, brown adipose tissue (BAT), white adipose tissue (WAT) and vena cava. Summary of the methods used (e.g., proteomics, transcriptomics, or multiplexed immunoassays) and availability of specific tissues for each exerkine is included in **Table S1**. Analyses were conducted using MotrpacRatTraining6moData R package (version 2.0.0) in RStudio (version 4.2.2).

Significance was assessed using three consecutive methods. P-adjusted values were used to evaluate overall significance. If the P-adjusted value was significant (< 0.05), we used the sex-specific p-values to determine significance. In all cases, if the error bars crossed the zero value, significance was deemed negligible regardless of the p-value [12]. Given that the MoTrPaC dataset uses different multiplexed immunoassay kits including the Rat Cytokine/Chemokine 27- Plex Discovery Assay (rat-27magplex), MILLIPLEX Rat Metabolic Hormone Magnetic Bead Panel (rat-metabolic), MILLIPLEX Rat Myokine Magnetic Bead Panel (rat-myokine) and MILLIPLEX MAP Rat Pituitary Magnetic Bead Panel (rat-pituitary). In cases where two immunoassays were used to measure the same exerkine/marker, significance was considered only if both assays showed a p-value < 0.05 and followed the same time- and sex-specific patterns. Similarly, if three immunoassays were employed, at least two must have shown a p-value < 0.05 and followed the same time- and sex-specific patterns.

## Results

Due to the scarcity of comprehensive studies integrating the responsiveness of different tissues to identical exercise regimens, we investigated which tissues were most responsive to exerkine alterations. The 10 most exercise-responsive tissues (**Figure 1C**) were BAT (11 exerkines) > adrenal gland (7 exerkines) > WAT = lungs (6 exerkines) > blood cells = vastus lateralis = spleen (4 exerkines) > cortex = hippocampus = heart (3 exerkines). The hypothalamus, gastrocnemius and vena cava had no exerkine alterations. The combined tissue exercise-responsiveness patterns held true for most exerkines when stratified by males (**Figure 1D**) or females (**Figure 1E**). These results are summarized in **Table S2**. Most of these exercise-related alterations were related to transcriptional changes (**Figure 1F-H**). This could be attributed either to the exerkines being more responsive at the transcriptional level or to the predominance of RNA-based quantification methods for the majority of tissues and exerkines, while protein quantification using either proteomics or multiplexed immunoassays was scarcer (see **Table S1** for more details).

In addition, we systematically quantified the number of tissue-specific alterations for each exerkine to uncover patterns of differential responsiveness. The 10 most exercise-responsive exerkines (**Figure 1F**) were fractalkine/CX3CL1 (8 tissues) > PSAP (7 tissues) > Cathepsin B (6 tissues) > Clusterin = FNDC (5 tissues) > PF4 = SPARC (4 tissues) > BDNF = IL-10 = Gpld1 (3 tissues). The combined exerkine-tissue alteration patterns held true for most exerkines when stratified by males (**Figure 1G**) or females (**Figure 1H**). Six exerkines (ADIPOQ, ANGPT1, APLN, LIF, TGFβ1, and TGFβ2) were altered in only two tissues, while four exerkines (GDF15, IL-15, METRNL, and myostatin) were altered only in one tissue. Seven exerkines (FGF21, IL-6, IL-7, IL-13, Musclin, SDC4, oxytocin) were not altered across any tissues. These results including the sex-specific % responsiveness for each tissue and exerkine are summarized in **Table S3**.

To elucidate any time-dependent, intensity-related, or sex-dimorphic effects for each tissue and exerkine, we stratified our responsiveness analysis based on whether the alteration occurred at 1 or 2 weeks (short-term, low-intensity), 4 weeks (mid-term, moderate-intensity), or 8 weeks (long-term, high-intensity) for both males and females (see **Figure 1A** for details). For short-term low-intensity exercise (**Figure S1A-C**) males were more responsive than females in tissues such as BAT (6 vs. 2 alterations), WAT (6 vs. 2 alterations), spleen (4 vs. 1 alterations), and less responsive in tissues such as the vastus lateralis (1 vs. 3 alterations). For mid-term moderate-intensity exercise (**Figure S1D-F**), males were more responsive than females in tissues such as the lungs (6 vs. 3 alterations), BAT (5 vs. 2 alterations), WAT (5 vs. 3 alterations), spleen and plasma (2 vs. 0 alterations), but were substantially less responsive in the adrenal gland (2 vs. 6 alterations). For long-term high-intensity exercise (**Figure S1G-I**), similar patterns occurred with the exception that WAT has become equally responsive in males and females. Across all time points, most tissues exhibited either equal or decreased responsiveness, with the exception of BAT, which demonstrated the highest responsiveness during long-term high-intensity exercise. Summary of time- and intensity-related changes for exerkines can be seen in Figures S2A-I. These results are summarized in **Table S4**.

### Exerkine-specific results

#### APLN

Though apelin was first purified and discovered in 1998 from bovine stomach tissue [13], its role as an exerkine is known to be mostly related to skeletal muscle of exercised humans and rodents [14; 15]. Several studies have demonstrated that apelin is capable of reversing age-related degenerative conditions such as a sarcopenia [15; 16]. In agreement with previous reports showing that exercise increases apelin transcription in muscles of humans and rodents, exercise increased *Apln* gene expression in the vastus lateralis of females (p < 0.0001), but not males (p = 0.22), across all 8 weeks (**Figure S3A**). In the lungs, *Apln* gene expression initially decreased in males (p = 0.016) and females (p = 0.027) at 1 week and gradually increased by 8 weeks (**Figure S3B**), although the adjusted p-value did not quite reach significance. Apelin has been shown to have a well-documented protective role against lungs respiratory diseases [17], but its levels were unchanged in the lungs (**Figure S4A**). APLN levels were unchanged in the cortex (**Figure S4B**). *Apln* gene expression was unchanged in the brain (cortex, hippocampus, hypothalamus), skeletal muscle (gastrocnemius), heart, kidney, adrenal gland, colon, spleen, testes, ovaries, small intestine, BAT, WAT and vena cava (**Figure S4C-Q**).

#### ADIPOQ

Adiponectin was first identified in 1995 as a primarily adipose tissue protein that significantly impacts metabolic processes such as glucose regulation and fatty acid breakdown [18]. Several studies have shown that it can have neurotrophic and neuroprotective effects, and also plays significant roles in reducing inflammation, enhancing insulin sensitivity, and mitigating atherogenic processes [1; 19; 20]. Several studies have shown that exercise increases adiponectin levels in the circulation of humans [21–23], whereas rodent studies show conflicting results [24–26].

Exercise decreased ADIPOQ plasma levels only in males (p < 0.001) at 1 and 8 weeks (**Figure S5A**), in contrast to a study showing that male rodents with free-access to voluntarily wheel running for 4-weeks, first show a decrease in adiponectin plasma levels at 1 week then gradually increase for the next three weeks [24]. Adiponectin levels and gene expression were unchanged in the gastrocnemius and vastus lateralis (**Figure S5B-E**), BAT (**Figure S5F-G**) or WAT (**Figure S5H-J**). *Adipoq* expression was unchanged in the brain (**Figure S5K-M**) despite reports in the literature [20; 27–29]. *Adipoq* gene expression in the adrenal gland showed a sex-dimorphic response with males (p < 0.0001) increasing and females (p < 0.0001) decreasing across all 8 weeks (**Figure S5N**). ADIPOQ levels were unchanged in the heart, lungs and kidney (**Figure S6A-C**). *Adipoq* gene expression was unchanged in the heart, kidney, colon, spleen, testes, ovaries, lungs, small intestine and vena cava (**Figure S6D-L**).

#### METRNL

METRNL has important roles in regulating immune responses and metabolism, and is expressed in various tissues including skeletal muscle, BAT and WAT, and is known to be exercise-responsive and cold-sensitive [30]. One of its main functions is the facilitation of BAT to WAT conversion and increasing adipose tissue thermogenesis in male rodents [31]. Indeed, exercise upregulated *Metrnl* gene expression in BAT only for males (p < 0.0001) at 2, 4 and 8 weeks of exercise (**Figure S7A**). METRNL levels and *Metrnl* gene expression were unchanged in WAT (**Figure S7B, Figure S8A**). *Metrnl* gene expression was unchanged in blood cells, the brain (cortex, hippocampus, hypothalamus), skeletal muscle (gastrocnemius and vastus lateralis), heart, kidney, adrenal gland, colon, spleen, testes, ovaries, lungs, small intestine and vena cava (**Figure S8B-R**).

#### FGF21

FGF21 is a hormone predominantly produced by the liver with profound effects on glucose and lipid metabolism. It plays a role in the body’s response to stress and starvation by promoting lipolysis, ketogenesis, and insulin sensitivity. FGF21 can convert WAT to BAT, and acts on the brain to regulate energy expenditure, food intake, and body weight [32]. It is also modulated by the exerkine adiponectin [33]. Acute exercise has been shown to increase FGF21 levels in the circulation of humans and in the circulation and liver, but not skeletal muscle or WAT, of rodents [34]. FGF21 levels were unchanged in the plasma, brain (cortex, hippocampus), skeletal muscle (gastrocnemius, vastus lateralis), heart, testes, ovaries, BAT and WAT (**Figure S9A-J)**. FGF21 levels and gene expression were not assessed in the liver.

#### GDF15

GDF15 is a stress-responsive cytokine from the TGF-β superfamily, known for its roles in regulating inflammation and metabolism. Primarily produced during cellular stress and disease states, GDF15 acts on the brain to regulate energy metabolism and appetite. It also influences WAT, promoting its conversion to BAT and stimulating thermogenesis [35; 36]. Two studies have shown that GDF15 plasma levels are immediately elevated after acute exercise in both male humans and rodents [37; 38]. Additionally, in rodents [38], this response extends to *Gdf15* gene expression in the liver, heart, and skeletal muscle (soleus). However, these effects were not observed when the exercise regimen utilized free-access to wheels, suggesting that the specific conditions of forced treadmill exercise are necessary to induce these changes in GDF15 levels and *Gdf15* gene expression [37; 38]. For the first time, we report that exercise decreased *Gdf15* gene expression only in males (p < 0.0001) at 1 and 2 weeks and increased its expression at 4 and 8 weeks in BAT (**Figure S10A**). *Gdf15* gene expression was unchanged in blood cells, the brain (hippocampus, hypothalamus), heart, kidney, adrenal gland, colon, spleen, ovaries, lungs, small intestine, liver, WAT and vena cava (**Figure S10B-O**).

#### FNDC5/Irisin

FNDC5/irisin has become a very popular exerkine due to its hypothetical effects on metabolism and overall health. Irisin is a peptide hormone that is cleaved from the type I membrane protein FNDC5, and irisin is thought to be predominantly produced by muscle tissue during exercise [39]. Male and female mice with free access to a running wheel for three weeks, and male humans undergoing 10 weeks of aerobic training, showed increased *FNDC5/Fndc5* gene expression in the quadriceps, alongside higher plasma irisin levels [40]. Since then, several studies have reported irisin elevations in response to exercise in humans and rodents [41–43], although some of these findings remain highly debatable [44; 45].

Herein, exercise almost decreased (p = 0.066) FNDC5 levels in both males and females in the vastus lateralis (**Figure 2A**). In WAT, exercise also appeared to decrease (p = 0.052) FNDC5 levels in males at 4 weeks but increase it at 8 weeks, while in females, the levels remained stable across all 8 weeks (**Figure 2B)**. These results contradict previous findings showing that FNDC5 levels are increased in skeletal muscle and WAT in response to exercise [46]. Additionally, *Fndc5* gene expression decreased in males (p < 0.0001) for the first 4 weeks, but then increased at 8 weeks, while in females (p = 0.039), it increased across all 8 weeks in the vastus lateralis (**Figure 2C**). *Fndc5* gene expression in the adrenal gland also showed a sex-dimorphic response to exercise, increasing in males (p < 0.001) at 1, 2 and 8 weeks but decreasing in females (p < 0.0001) across all 8 weeks (**Figure 2D**). In the liver, *Fndc5* gene expression increased only in males (p = 0.0004) at 2 and 4 weeks, but decreased 8 weeks (**Figure 2E**). In BAT, *Fndc5* gene expression decreased at 8 weeks in both males and females, but this was only significant (p < 0.001) for females (**Figure 2F**). In WAT, exercise showed a sex-dimorphic response; it decreased *Fndc5* gene expression in males (p = 0.0007) across all 8 weeks (**Figure 2G),** while in females, it appeared to increase it across all 8 weeks, although this was not significant (p = 0.12).

**Figure 2.**
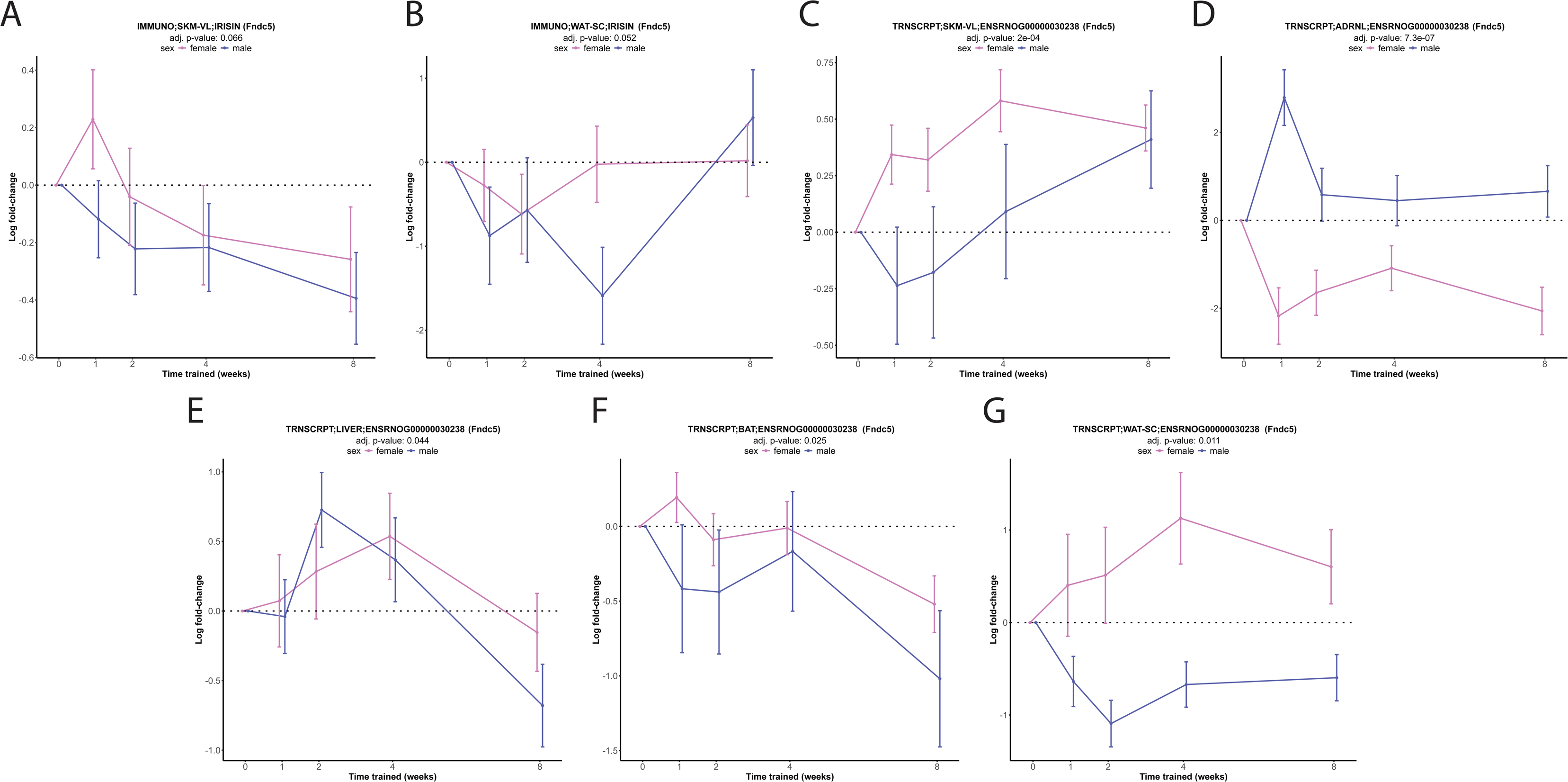
Analysis of fibronectin type III domain-containing protein 5 (Fndc5)/irisin levels and gene expression of 6-month treadmill-exercised rats across all 8 weeks of training. FNDC5 levels in vastus lateralis (SKM-VL) and white adipose tissue (WAT) (**A-B)**. *Fndc5* gene expression in the vastus lateralis (SKM-VL), adrenal gland (ADRNL), liver, brown adipose tissue (BAT) and WAT (**A-G**).

FNDC5 levels were unchanged in plasma, brain (cortex, hippocampus), skeletal muscle (gastrocnemius), heart, testes, ovaries and BAT (**Figure S11A-H**). It has been reported that several ELISA kits could not reliably distinguish irisin plasma concentrations between *Fndc5* knockout and wild-type mice [47], but Luminex bead-based technology (used in the MoTrPaC study) was not employed in these assessments. *Fndc5* gene expression was unchanged in blood cells, the brain (cortex, hippocampus, hypothalamus), skeletal muscle (gastrocnemius, vastus lateralis), heart, kidney, colon, spleen, testes, ovaries, lungs, small intestine and vena cava (**Figure S11I-Y**). Given that FNDC5/Irisin is often thought of as a muscle- or adipocyte-originating brain-targeting exerkine [39], the non-changing levels and expression in these tissues (skeletal muscle, WAT and brain) contradicts this notion and supports some of the opposing views [44; 45].

#### Cathepsin B

CTSB is a lysosomal cysteine protease important for protein turnover within cells. There has been debate in the field on whether cathepsin B is an exerkine with studies in young male humans showing that acute exercise does not increase CTSB plasma levels [48], and other studies with young male/female humans, rhesus monkey and voluntary wheel accessing mice showing that chronic exercise increases CTSB plasma levels [49; 50]. Moreover, in young male and female mice given access to a voluntary wheel for 30 days, *Ctsb* gene expression increased in the gastrocnemius skeletal muscle and decreased in the spleen, while it remained unchanged in the soleus, WAT, liver and frontal cortex [50].

Exercise decreased CTSB WAT levels only in males (p = 0.0088) at 1, 2 and 4 weeks before returning to baseline at 8 weeks (**Figure 3A**). *Ctsb* gene expression changed in different tissues and across various time points, showing sex-specific patterns. In males, exercise altered *Ctsb* gene expression in blood cells across all 8 weeks (**Figure 3B**, p < 0.0001), spleen at 2 and 8 weeks (**Figure 3D**, p = 0.0071), lungs across all 8 weeks (**Figure 3E**, p = 0.035) and BAT at 8 weeks (**Figure 3F**, p = 0.0002). For females, exercise decreased *Ctsb* gene expression in the colon at 1, 2 and 8 weeks (**Figure 3C**, p = 0.0008), increased *Ctsb* gene expression in the spleen across at 1, 2 and 8 weeks (**Figure 3D**, p = 0.025), and decreased *Ctsb* gene expression in the lungs during the first four weeks (**Figure 3E**, p = 0.0019). Exercise did not induce post-translational modifications of CTSB including acetylation in the liver (**Figure S12A-B**) nor its phosphorylation in the gastrocnemius heart and kidney (**Figure S12C-H**).

**Figure 3.**
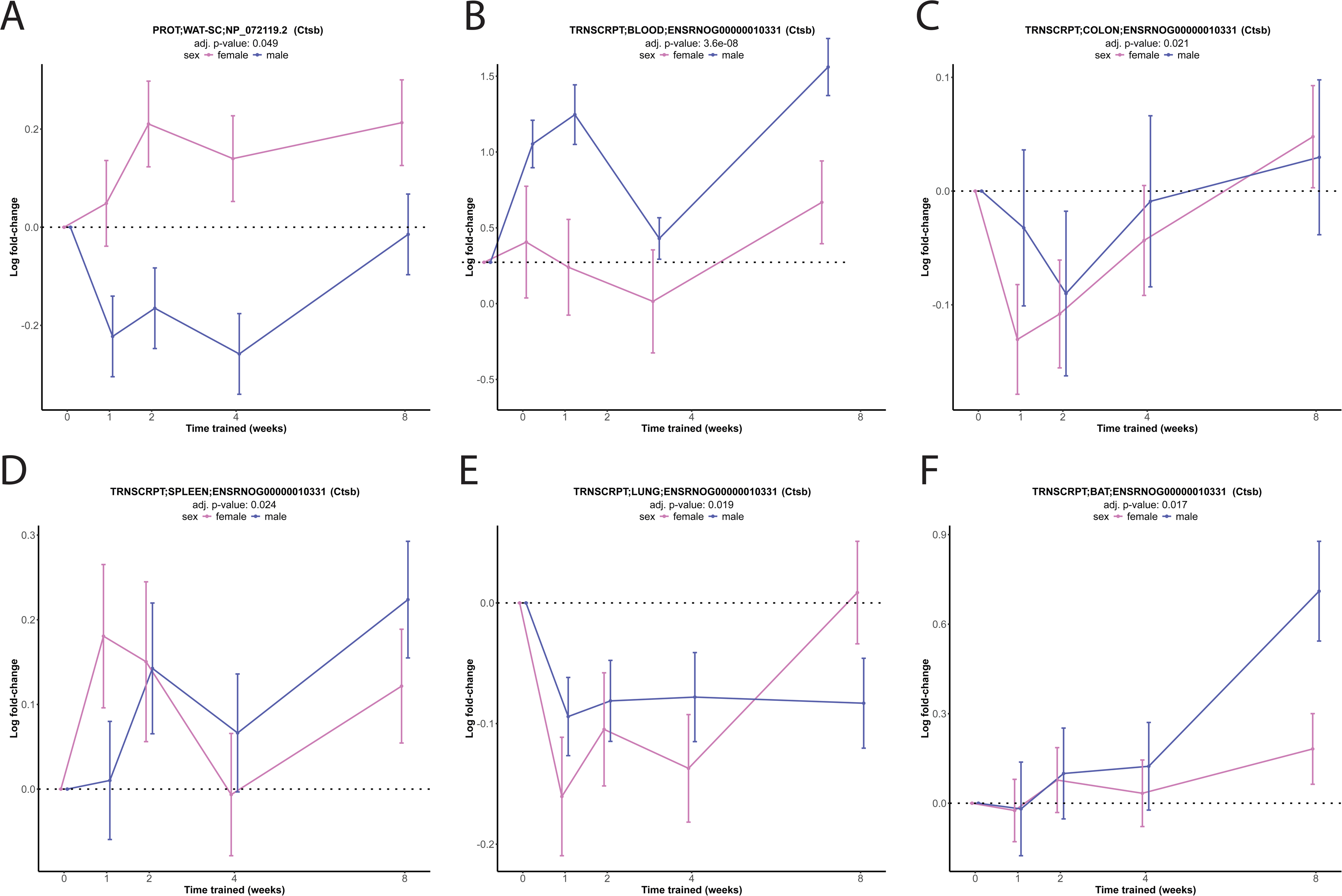
Analysis of cathepsin B (CTSB) levels and gene expression of 6-month treadmill-exercised rats across all 8 weeks of training. CTSB levels in white adipose tissue (WAT) (**A)**. *Ctsb* gene expression in blood cells, colon, spleen, lungs and brown adipose tissue (BAT) (**B-F**).

Despite reports speculatively claiming that CTSB is a muscle-originating brain-targeting exerkine without sufficient evidence, results herein show that exercise did not alter CTSB levels in the cortex, gastrocnemius (**Figure S12I-J**) or *Ctsb* gene expression in the brain (cortex, hippocampus, hypothalamus) or skeletal muscle (gastrocnemius, vastus lateralis) (**Figure S13A- E**). Collectively, these suggest that CTSB is not a brain-targeting exerkine, and is in alignment with the fact that only very few studies exist to support this claim despite the first report in 2016 [50]. CTSB levels were unchanged in the heart, kidney, lungs and liver (**Figure S12K-O**). *Ctsb* gene expression was unchanged in the heart, kidney, adrenal gland, testes, ovaries, liver, WAT and vena cava (**Figure S13F-M**).

#### PSAP

PSAP is a multifunctional lysosomal protein that was first discovered in 1988 [51]. PSAP serves as a precursor to saposins A-D, important molecules for the proper degradation of lipids within lysosomes. PSAP plays an essential role in lipid metabolism and neural development. Deficiencies or mutations in the PSAP gene are linked to several lipid storage and neurodegenerative disorders. PSAP also has protective roles in the nervous system and has been studied for its potential therapeutic benefits in neurodegenerative conditions [52].

Although PSAP has not yet been identified as an exerkine, its levels have been shown to increase in the skeletal muscle, adipose tissue, and their extracellular fluids of PGC1-α overexpressing mice and cell cultures, as well as in response to cold exposure in adipose tissue. However, the same study found that acute treadmill exercise did not alter PSAP levels in the skeletal muscle and its extracellular fluid of young male mice, as evidenced by its absence in the significant ‘hits’ of the proteomics analysis [10]. Additionally, a recent aptamer-based proteomics study showed that acute exercise decreases PSAP plasma levels in older male humans [53]. As such, the presumption that PSAP is an exerkine or exercise-responsive molecule remains largely speculative.

Interestingly, exercise altered PSAP levels in WAT in a sex-dimorphic manner; levels decreased for the first 4 weeks in males (p = 0.021) before returning to baseline at 8 weeks, while consistently increasing in females (p = 0.0106) across all 8 weeks (**Figure 4A**). This was not extended to *Psap* gene expression in WAT (**Figure S14A**), but similar significant trends were seen for *Psapl1* gene expression in WAT for males (p = 0.043) at 2 and 4 weeks (**Figure 4B**). Interestingly, the WAT male data (**Figure 4A-B, Figure S14A**) contradicts the only study [10] demonstrating that cold exposure increases PSAP levels and gene expression in WAT and PSAP levels in WAT’s extracellular fluid of young male mice. However, the study did not assess PSAP levels in WAT post-exercise or in females. Additionally, *Psap* gene expression decreased in the vastus lateralis in males (p = 0.033) at 8 weeks, while in females (p < 0.0001), it decreased across all 8 weeks (**Figure 4C**). PSAP levels and gene expression were unchanged in the gastrocnemius (**Figure S14B-C**), contradicting the results with PGC1-α [10] and transcription factor E-B (TFEB) [11] muscle-overexpressing mice claiming that PSAP is a metabolic-responsive myokine. PSAP levels were unchanged in the cortex (**Figure S14D-E**). *Psap* gene expression was unchanged in the cortex, hippocampus and hypothalamus (**Figure S14F-H)**, suggesting that it is not a brain-targeting exerkine.

**Figure 4.**
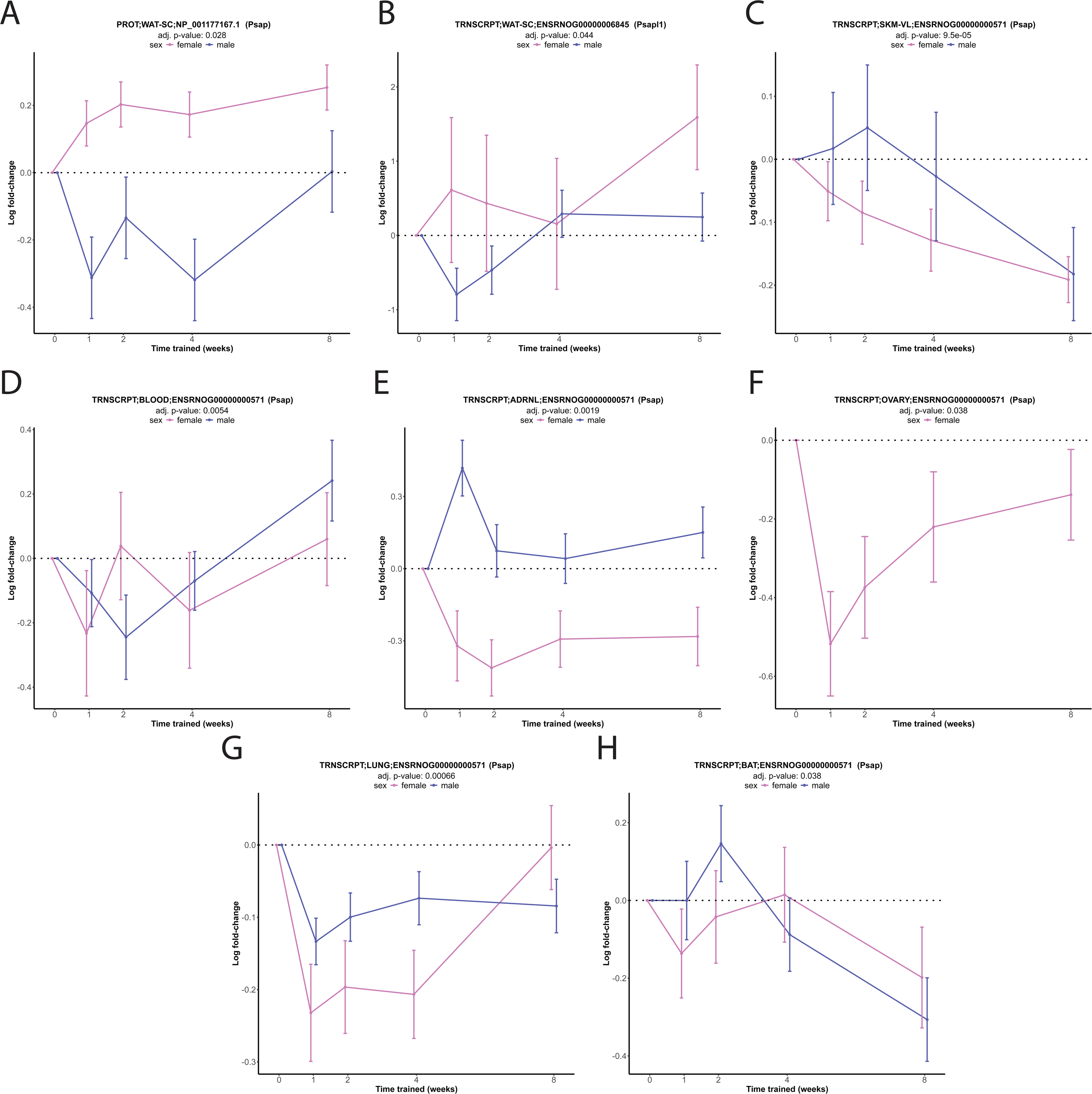
Analysis of prosaposin (Psap) levels and gene expression of 6-month treadmill-exercised rats across all 8 weeks of training. PSAP levels in white adipose tissue (WAT) (**A)**. *Psap* like 1 gene expression in WAT (**B**). *Psap* gene expression in the vastus lateralis (SKM- VL), blood cells, adrenal gland (ADRNL), ovaries, lungs and brown adipose tissue (BAT) (**C- H**).

In blood cells, *Psap* gene expression decreased only in males (p < 0.0001) at 2 and 4 weeks, but increased at 8 weeks (**Figure 4D**). In the adrenal gland, *Psap* gene expression increased in males (p = 0.0021) at 1 and 8 weeks, while in females (p = 0.0059), it decreased across all 8 weeks (**Figure 4E**). In the ovaries, *Psap* gene expression decreased across all 8 weeks (**Figure 4F**). In the lungs, *Psap* gene expression decreased in males (p = 0.001) across all 8 weeks, while in females (p = 0.0006), it decreased for the first 4 weeks before returning to baseline (**Figure 4G**). In BAT, *Psap* gene expression increased and decreased only in males (p = 0.0005) at 2 and 8 weeks, respectively (**Figure 4H**). PSAP acetylation was unchanged in the heart and kidney (**Figure S14I-L**). PSAP phosphorylation was unchanged in the liver (**Figure S14M-N**). PSAP levels were unchanged in the heart, kidney, lungs and liver (**Figure S14O-S**). *Psap* gene expression was unchanged in the heart, kidney, spleen, testes, small intestine and vena cava (**Figure S14T-ZA**).

#### IL-6

IL-6 a cytokine that plays a pleiotropic role across immune responses, inflammation, hematopoiesis, maintenance of skeletal muscle and bone mass, brain activity and overall metabolism [54]. It was first identified in the 1980s as a factor responsible for inducing the production of acute-phase proteins during inflammation and for stimulating B cell differentiation. Over the years, IL-6 has been shown to have both pro- and anti-inflammatory effects [55]. IL-6 was identified in 2000 as a skeletal muscle exerkine in male humans following an acute exercise bout of one-legged knee extensions [56]. Subsequent studies have shown that IL-6 plasma levels are elevated in response to both short- and long-term exercise in humans and rodents [1; 57; 58]. Surprisingly, IL-6 levels were unchanged in the plasma, brain (cortex, hippocampus), skeletal muscle (gastrocnemius, vastus lateralis), heart, kidney, adrenal gland, colon, spleen, testes, ovaries, lungs, liver, BAT and WAT (**Figure S15A-O**).

#### IL-7

IL-7 is a cytokine primarily involved in inflammatory responses and is thought of as an exerkine. Its role during physical activity pertains to modulating inflammation and aiding in muscle repair [59; 60]. The relationship between exercise and IL-7 levels in both humans and rodents continues to be an active area of research, focusing on how it might influence recovery and immune responses post-exercise. Exercise did not change *Il-7* gene expression in blood cells, the kidney, colon, spleen, ovaries, lungs, small intestine, liver, BAT, WAT and vena cava (**Figure S16A-K**).

#### IL-10

IL-10 is an anti-inflammatory cytokine and exerkine, shown extensively to be elevated during exercise in humans [61], and to a lesser extent in rodents [62]. Its production is often stimulated by IL-6, highlighting a regulatory mechanism where IL-6 promotes IL-10 release to mitigate inflammation and aid in post-exercise recovery [63]. Exercise appeared to decrease IL-10 plasma levels in females across all 8 weeks, while in males, levels gradually decreased from weeks 1 to 4, then suddenly increased at 8 weeks, indicating a sex-dimorphic response (**Figure S17A**), although this was not significant. In the cortex, IL-10 levels decreased in females (p = 0.036) from 1 to 4 weeks, and returned to baseline at 8 weeks, but in males (p = 0.056), they were unchanged (**Figure S17B**). IL-10 levels were unchanged in the hippocampus (**Figure S17C**). Despite IL-10 being thought of as a skeletal muscle exerkine [64], IL-10 levels were unchanged in the gastrocnemius and vastus lateralis (**Figure S17D-E**). In the heart, IL-10 levels increased in females (p = 0.0066) across all 8 weeks (**Figure S17F**). In the spleen, IL-10 levels decreased in males (p = 0.008) at 1, 4 and 8 weeks (**Figure S17G**). IL-10 levels were unchanged in the adrenal gland, kidney, testes, ovaries, lungs, small intestine, liver, BAT and WAT (**Figure S17H-P**).

#### IL-13

IL-13 is a T helper 2 cell cytokine, known for its role in the anti-inflammatory response and macrophage polarization within WAT [65]. Beyond IL-13’s immunological functions, it has been shown to be an exerkine. It increases in the circulation in male/female humans following 4 weeks of aerobic exercise and in young male/female rodents following 4 weeks of treadmill exercise. Additionally, deficiencies in IL-13 lead to reduced exercise capacity and diminished metabolic benefits in rodents [66]. Exercise did not alter IL-13 levels in the plasma, brain (cortex, hippocampus), skeletal muscle (gastrocnemius, vastus lateralis), heart, kidney, adrenal gland, colon, spleen, testes, ovaries, lungs, small intestine, liver, BAT and WAT (**Figure S18A- P**).

#### IL-15

IL-15 is a cytokine responsible for immune system regulation, enhancing the development and function of natural killer, T and B cells. Several studies reported that IL-15 circulation levels are not altered following acute [67–70] or chronic [71; 72] exercise in humans, while others have reported such elevation only after acute exercise [73; 74] or chronic exercise in females only [75]. Rodent studies reported similar contradictory findings [76; 77]. IL-15 have been shown to induce lipolysis and reduce adipose tissue mass [78–80]. Interestingly, exercise decreased IL-15 WAT levels only in males (p = 0.004) at 1 and 4 weeks, but increased it at 8 weeks (**Figure S19A**). IL-15 levels were unchanged in the plasma, brain (cortex, hippocampus), skeletal muscle (gastrocnemius, vastus lateralis), heart, testes, ovaries and BAT **(Figure S19B-J**).

#### Musclin

Musclin/osteocrin was first identified as a skeletal muscle-originating factor in 2004 [81]. Musclin levels and gene expression have also been shown to be elevated in the skeletal muscle of young male/female rodents subjected to 5 days of treadmill exercise, with musclin levels additionally increased in the circulation [82]. Similarly, acute exercise increased musclin levels in the circulation of young inactive males during and up to 120 minutes after the exercise bout [83]. In this study, although exercise did not significantly alter musclin levels or gene expression across all assessed tissues, there was a trend toward elevated levels in females for plasma across all 8 weeks (**Figure S20A**), gastrocnemius from 1 to 4 weeks (**Figure S20B**) and vastus lateralis from 1 to 2 weeks (**Figure S20C**) quantified using an immunoassay. Musclin levels were unchanged in the brain (cortex, hippocampus), heart, testes, ovaries, BAT and WAT (**Figure S20D-J**). Musclin phosphorylation and levels were unchanged in skeletal muscle (**Figure S20K- M)** when assessed with proteomics. Musclin gene expression was unchanged in skeletal muscle (gastrocnemius, vastus lateralis), BAT, WAT and vena cava (**Figure S20N-R**).

#### Myostatin/GDF8

Myostatin, also known as growth differentiation factor 8 (GDF-8), is a member of the TGFβ protein family that functions as a natural brake on muscle growth to prevent excessive hypertrophy. Released during exercise, myostatin levels typically decrease to promote muscle growth, yet a slight initial release can occur as a regulatory mechanism during intense physical activity, signaling the body to limit muscle expansion to safe levels [1; 84]. Previous studies have shown that myostatin acts as a negative regulator of WAT to prevent browning and thermogenesis, leading to less energy expenditure and higher risk of weight gain [85–87]. Exercise decreased MSTN WAT levels in males (p = 0.0094) at 1 and 8 weeks, suggesting enhanced metabolic activity and energy expenditure. In contrast, exercise increased MSTN WAT levels in females (p = 0.014) for the first four weeks. (**Figure S21A**). However, this was not replicated in the immunoassay analysis (**Figure S21B**). MSTN levels were unchanged in the gastrocnemius or vastus lateralis (**Figure S21C-E**). MSTN levels were unchanged in the plasma, brain (hippocampus, hypothalamus), heart, testes, ovaries and BAT (**Figure S21F-L**). *Mstn* gene expression was unchanged in brain (cortex, hippocampus, hypothalamus), skeletal muscle (gastrocnemius, vastus lateralis), spleen, BAT, WAT and vena cava (**Figure S21M-U**).

#### LIF

LIF is a pleiotropic cytokine belonging to the IL-6 family, known for its wide-ranging roles in cellular growth, differentiation, and survival. It is involved in various physiological processes, including immune regulation, neuronal development, and muscle repair. LIF is produced in response to stress or injury, such as during exercise, and can promote regeneration by activating signaling pathways like JAK/STAT in muscle and other tissues [88]. Its transient increase during exercise suggests it plays a role in tissue adaptation and repair, although its effects depend on factors like exercise type, intensity, and duration [89; 90]. Though LIF is well-known to have protective effects in the heart, including stimulating repair and preventing cellular damage [91], exercise decreased LIF heart levels only in males (p = 0.001) from 1 to 4 weeks (**Figure S22A**). We are unaware of studies investigating LIF levels in the heart after exercise. In one study, chronic exercise increased LIF expression and prevented myocardial-infarction-induced skeletal muscle atrophy in rodents [92]. Interestingly, exercise also decreased LIF BAT levels only in males (p = 0.0026) at 1 and 4 weeks, but increased it at 8 weeks (**Figure S22B**). LIF levels were unchanged in the brain (cortex, hippocampus), skeletal muscle (gastrocnemius, vastus lateralis), testes, ovaries and WAT (**Figure S22C-J**).

#### SPARC

SPARC is involved in various physiological processes including wound healing, bone mineralization, angiogenesis and extracellular matrix remodeling. SPARC has been shown to be secreted in response to exercise in both humans and rodents [93; 94]. Exercise increased SPARC levels in males (p = 0.012) at 4 and 8 weeks and dynamically altered the levels in females (p = 0.0028), increasing at 1 and 4 weeks and decreasing at 2 weeks in the cortex (**Figure S23A**). In the hippocampus, SPARC levels increased in males (p = 0.0017) at 4 weeks, and in females (p = 0.0028) at 1, 4 and 8 weeks (**Figure S23B**). In the skeletal muscle (gastrocnemius and vastus lateralis), SPARC levels decreased in males at 8 weeks, but this was only significant for the vastus lateralis (p = 0.0006), while in females, SPARC levels initially increased from 1 to 4 weeks (**Figure S23C-D**). In testes, SPARC levels increased at 1 and 2 weeks (**Figure S23E**). SPARC levels were unchanged in the heart, BAT, WAT and ovaries (**Figure S23F-H**).

#### SDC4

SDC4 is a transmembrane heparan sulfate proteoglycan that plays a role in regulating cellular processes such as migration, proliferation, and differentiation through its interaction with extracellular matrix proteins. SDC4 may contribute to the remodeling of skeletal muscle by mediating signaling pathways that promote muscle repair and growth [95]. Exercise has been shown to increase SDC4 circulation levels in young male humans [96] and rodents [97]. It has also been suggested that SDC4 is a hepatokine [97]. Exercise did not alter SDC4 levels in the cortex, kidney, lungs, liver and WAT (**Figure S24A-E**). Post-translational modifications were unchanged in the heart (phosphorylation) and liver (phosphorylation or ubiquination) (**Figure S24F-I**). *Sdc4* gene expression was unchanged in blood cells, the brain (cortex, hippocampus, hypothalamus), skeletal muscle (gastrocnemius, vastus lateralis), heart, kidney, adrenal gland, colon, spleen, testes, ovaries, lungs, BAT, WAT and vena cava (**Figure S24J-ZA**).

#### TGFβ1and TGFβ2

TGFβ1 and TGFβ2 are part of the TGFβ superfamily of cytokines, which are key regulators in cellular growth, proliferation, differentiation, and apoptosis. TGFβ1 is widely known for its role in immune system regulation, fibrosis, and angiogenesis, and is generally considered a major fibrotic factor in various pathological conditions. On the other hand, TGFβ2 has similar biological functions but is often found in specific tissues such as the brain, heart, and eyes, contributing to embryonic development and the maintenance of tissue homeostasis [98]. They are both known to be elevated in response to acute and/or chronic exercise in humans and rodents [99–102]. TGFβ1 has important roles in the skeletal muscle [99] and exercise has been shown to increase its expression there in humans and rodents [100; 101]. In contrast, TGFβ2 is thought to be an exerkine originating from WAT [102]

Exercise did not alter TGFβ1 levels in the cortex, heart, kidney, lungs, liver and WAT (**Figure S25A-F).** Exercise increased *Tgf*β*1* gene expression in males (p < 0.001) at 4 and 8 weeks, and almost significantly (p = 0.061) decreased its expression in females at 2 and 4 weeks in BAT (**Figure S25G**). A similar trend was observed in WAT for males (p = 0.015) at 1, 2 and 4 weeks and females (p = 0.014) at 4 and 8 weeks (**Figure S25H**). Despite its known roles in skeletal muscle, *Tgf*β*1* gene expression was unchanged in skeletal muscle (gastrocnemius, vastus lateralis) (**Figure S25I-J**). *Tgfb1* gene expression was also unchanged in blood cells, the brain (cortex, hippocampus, hypothalamus), heart, kidney, adrenal gland, colon, spleen, testes, ovaries, lungs, small intestine, liver and vena cava (**Figure S25K-Y**).

Similarly, despite TGFβ2 known role as a WAT-originating exerkine, exercise did not alter its levels in WAT nor its gene expression in WAT and BAT (**Figure S26A-C**). TGFβ2 levels were also unchanged in the cortex, gastrocnemius, heart, and lungs (**Figure S26D-G**). In the colon, *Tgf*β*2* gene expression decreased in males (p = 0.0023) at 2 and 4 weeks, but increased at 8 weeks (**Figure S26H**). In the lungs, *Tgfb2* gene expression decreased in males at 2, 4 and 8 weeks (p < 0.0001), while in females (p < 0.0001), it initially increased at 2 weeks then decreased at 8 weeks (**Figure S26I**). *Tgf*β*2* gene expression was unchanged in blood cells, the brain (cortex, hippocampus, hypothalamus), skeletal muscle (gastrocnemius, vastus lateralis), heart, kidney, adrenal gland, spleen, testes, ovary, small intestine, liver and vena cava (**Figure S26J-X**).

#### ANGPT1

ANGPT1 is a growth factor essential for vascular stabilization and angiogenesis. It activates the Tie2 receptor on endothelial cells, enhancing vascular integrity and reducing permeability. It has been shown to increase in response to acute exercise in humans and rodents [103–105], with variables results in response to chronic exercise [104; 105]. Exercise did not alter ANGPT1 levels in the kidney, lungs and WAT (**Figure S27A-C**). Exercise decreased *Angpt1* gene expression in the ovaries consistently across all 8 weeks (**Figure S27D**). In the female (p = 0.0003) adrenal gland, exercise initially decreased ANGPT1 levels at 1 week, then increased them at 2 and 4 weeks (**Figure S27E**). *Angpt1* gene expression was unchanged in blood cells, the brain (cortex, hippocampus, hypothalamus), skeletal muscle (gastrocnemius, vastus lateralis), heart, kidney, colon, spleen, testes, lungs, small intestine, BAT, WAT and vena cava (**Figure S27F-V**).

#### Fractalkine/CX3CL1

Fractalkine, also known as CX3CL1, is a molecule that functions both as a chemokine and as an adhesion molecule. Exercise has been shown to variably affect fractalkine circulation levels in humans [106–108] and rodents [109]. As an exerkine, it has been shown to be released from skeletal muscle and act on immune cells [107]. Despite the limited number of studies investigating fractalkine levels and gene expression across tissues in response to exercise in rodents, in our analysis, exercise altered its protein levels in 8 tissues (cortex, hippocampus, heart, kidney, adrenal gland spleen, lungs and BAT), and its gene expression in 4 tissues (heart, adrenal gland, lungs and BAT).

In the cortex, exercise increased fractalkine levels in males at 8 weeks, whereas in females, levels fractalkine levels increased at 4 weeks before decreasing at 8 weeks (**Figure 5A-B**). In the hippocampus, fractalkine levels consistently decreased in males across all 8 weeks, whereas in females, they increased at 1 and 4 weeks (**Figure 5C-D**). In the heart, fractalkines levels decreased in males at 2 weeks, but increased at 8 weeks, whereas for females, levels increased only at 8 weeks (**Figure 5E-F**). Additionally, exercise altered fractalkine levels in various organs of males. In the kidney, fractalkine levels decreased consistently in males (p = 0.015) across all 8 weeks (**Figure 5G**). In the spleen, fractalkine levels increased only in males (p = 0.0017) at 2 weeks then decreased at 4 weeks (**Figure 5H**). In the lungs, fractalkine levels decreased only in males (p = 0.0043) at 2 and 4 weeks then increased at 8 weeks (**Figure 5I**). In BAT, fractalkine levels changed in a sex-dimorphic manner; in males, levels remained stable for the first 4 weeks and then suddenly increased at 8 weeks, whereas in females, levels consistently decreased across all 8 weeks (**Figure 5J-K**). Though exercise did not alter fractalkine levels in the WAT significantly, there was a trend toward decreased levels at 4 weeks for females, and increasing levels at 8 weeks for both males and females (**Figure 5L**).

**Figure 5.**
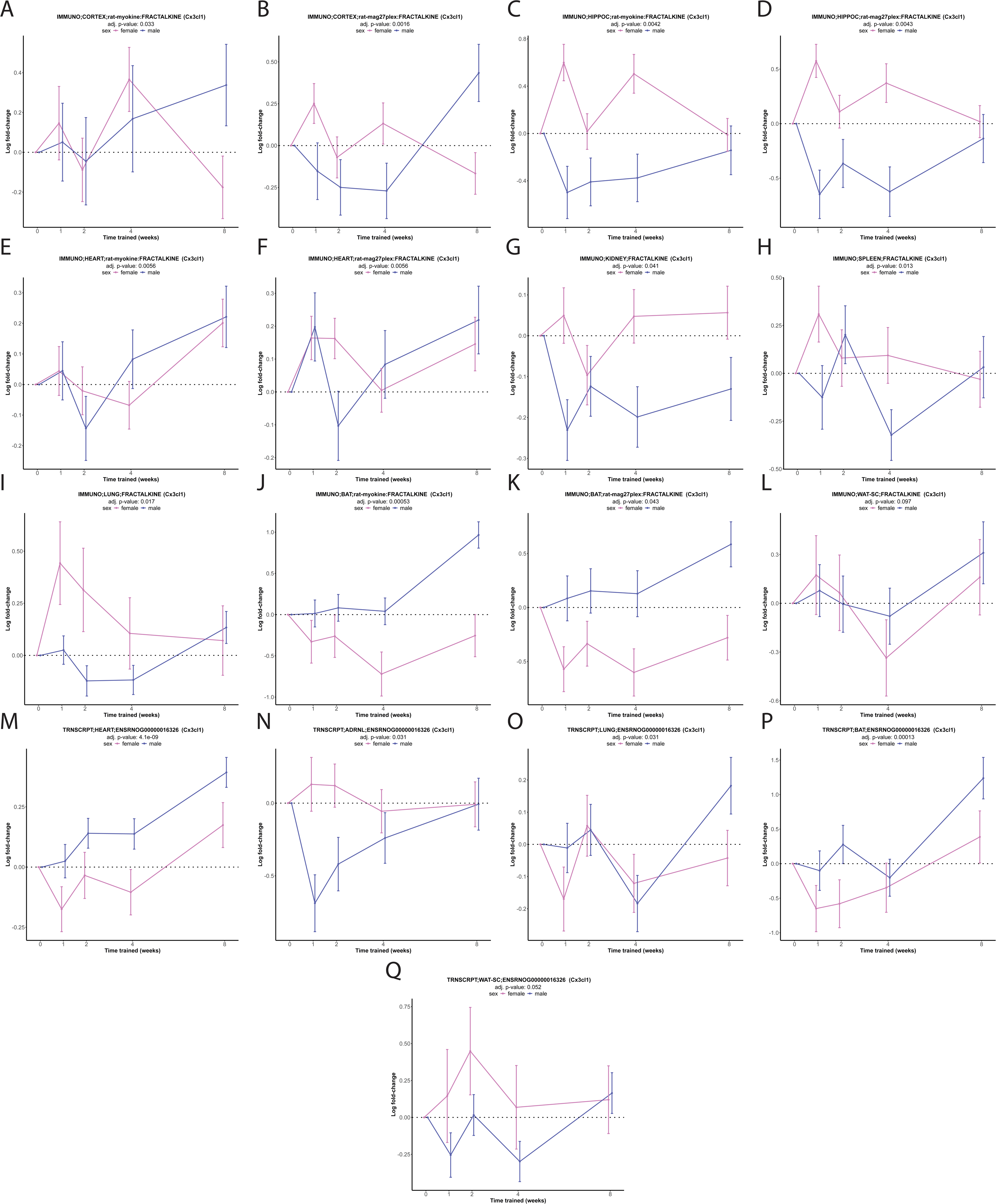
Analysis of fractalkine/C-X3-C motif chemokine ligand 1 (CX3CL1) levels and gene expression of 6-month treadmill-exercised rats across all 8 weeks of training. Fractalkine/CX3CL1 levels in the cortex, hippocampus (HIPPOC), heart, kidney, spleen, lungs, brown adipose tissue (BAT) and white adipose tissue (WAT) (**A-L**). *Cx3cl1* gene expression in the heart, adrenal gland (ADRNL), lungs, BAT and WAT (**M-Q**).

Exercise altered fractalkine gene expression in the heart for both males and females. In males (p < 0.0001), fractalkine gene expression increased from 2 to 8 weeks, while in females, it decreased at 1 and 4 weeks then increased at 8 weeks (**Figure 5M**). In the adrenal gland, fractalkine gene expression decreased only in males (p = 0.001) for the first 4 weeks before (**Figure 5N**). In the lungs, fractalkine gene expression differed by sex; in males (p = 0.0031), it remained stable for the first 2 weeks, decreased at 4 weeks, and then suddenly increased at 8 weeks, while in females (p = 0.048), it decreased at 1 and 8 weeks (**Figure 5O**). In BAT, fractalkine gene expression increased in males (p < 0.0001) at 8 weeks, whereas in females (p < 0.001), it decreased for the first 4 weeks before increasing at 8 weeks (**Figure 5P**). Although fractalkine gene expression in WAT was unchanged, there was a trend toward increasing levels for females and decreasing levels for males for the first 4 weeks (**Figure 5Q**).

Fractalkine levels were unchanged in the plasma, skeletal muscle (gastrocnemius, vastus lateralis), colon, testes, ovaries, small intestine, liver, cortex (including phosphorylation) and kidney (**Figure S28A-L)**. Fractalkine gene expression was unchanged in the brain (cortex, hippocampus, hypothalamus), skeletal muscle (gastrocnemius, vastus lateralis), kidney, colon, spleen, testes, ovaries, small intestine, liver and vena cava (**Figure S28M-Y**).

#### Oxytocin

Oxytocin is widely studied in the brain due its related effects to social bonding and stress relief. But it has also been shown to be important for muscle maintenance and regeneration [110]. Although the research on its role as an exerkine is limited, acute exercise has been shown to increase its circulating levels in male humans [111; 112]. In rodents, chronic exercise has been shown to increase circulating and brain levels of oxytocin only in females [113]. Exercise did not alter oxytocin levels in the cortex (**Figure S29A**) nor its gene expression in the cortex, hippocampus, hypothalamus and ovaries (**Figure S29B-E**).

#### BDNF

BDNF is a key molecule in the brain with well-documented effects in supporting neuronal survival, growth, differentiation of neurons, and learning and memory [114]. Acute exercise has been shown to increase BDNF levels in the circulation in male humans [48; 115], whereas results from chronic exercise are conflicting [116]. In rodents, acute and chronic exercise has been shown to not alter plasma BDNF levels [117; 118]. However, chronic exercise has been well-shown to upregulate BDNF brain levels, especially in the hippocampus [119]. This is supported by a recent meta-analysis showing that BDNF brain levels are upregulated in exercised male and female rodents [3; 120].

Exercise initially decreased BDNF plasma levels only in males (p < 0.001) at 1 week, but increased from 2 to 8 weeks (**Figure S30A-B**). Exercise appeared to decrease BDNF hippocampal levels in males at 1, 2 and 4 weeks, but this did not quite reach significance (p = 0.068) contrasting previous reports [116; 119; 120]. In females (p = 0.012), exercise consistently increased BDNF hippocampal levels across all 8 weeks (**Figure S30C**), but this effect was not replicated in the cortex (**Figure S30D**). Despite documented effects of BDNF in skeletal muscle [121–124], BDNF levels were unchanged in the gastrocnemius and vastus lateralis (**Figure S30E-F**). There have been several studies documenting the effect of exercise on browning of adipose tissue [125; 126]. Though studies evaluating BDNF levels in adipose tissue and browning are limited, one study showed that BDNF peripheral injection and acute exercise both elicit similar adipose tissue molecular alterations and browning [127]. Interestingly, BDNF BAT levels increased only in females (p = 0.0007) across all 8 weeks (**Figure S30G**), while in WAT the effect was opposite and almost reached significance (**Figure S30H**). BDNF levels were unchanged in the heart, adrenal gland, testes and ovaries (**Figure S31A-D**).

#### PF4

PF4 is a chemokine released by platelets that is traditionally associated with blood clotting but has recently been implicated in aging and neurological health [4; 8; 128; 129]. Additionally, PF4 levels have been shown to increase in the circulation following an acute bout of exercise in rodents [9; 130].

In blood cells, exercise increased *Pf4* gene expression only in males (p < 0.0001) at 8 weeks (**Figure 6A**), while in females, exercise consistently decreased *Pf4* gene expression across all 8 weeks, almost reaching significance (p = 0.076). Similarly, exercise decreased *Pf4* gene expression only in males (p < 0.0001) at 1 and 2 weeks before increasing it at 8 weeks in the spleen (**Figure 6B**). In the lungs, *Pf4* gene expression decreased only in males (p < 0.0001) across all 8 weeks (**Figure 6C**). In the small intestine, *Pf4* gene expression decreased only in females (p < 0.0001) at 1, 2 and 4 weeks (**Figure 6D**). PF4 levels were unchanged in the heart, kidney, liver, gastrocnemius, lungs and WAT (**Figure S32A-G**). *Pf4* gene expression was unchanged in WAT, gastrocnemius, vastus lateralis, heart, colon, ovaries, liver, BAT and vena cava (**Figure S32H-P**). Peripheral exercise-induced PF4 release has been shown to confer neuroprotective benefits and offer disease-modifying therapeutic potential as a brain-targeting exerkine [8; 9]. Unfortunately, PF4 levels and *Pf4* gene expression were not assessed in the cortex, hippocampus and hypothalamus in the MoTrPaC rat 6-month dataset.

**Figure 6.**
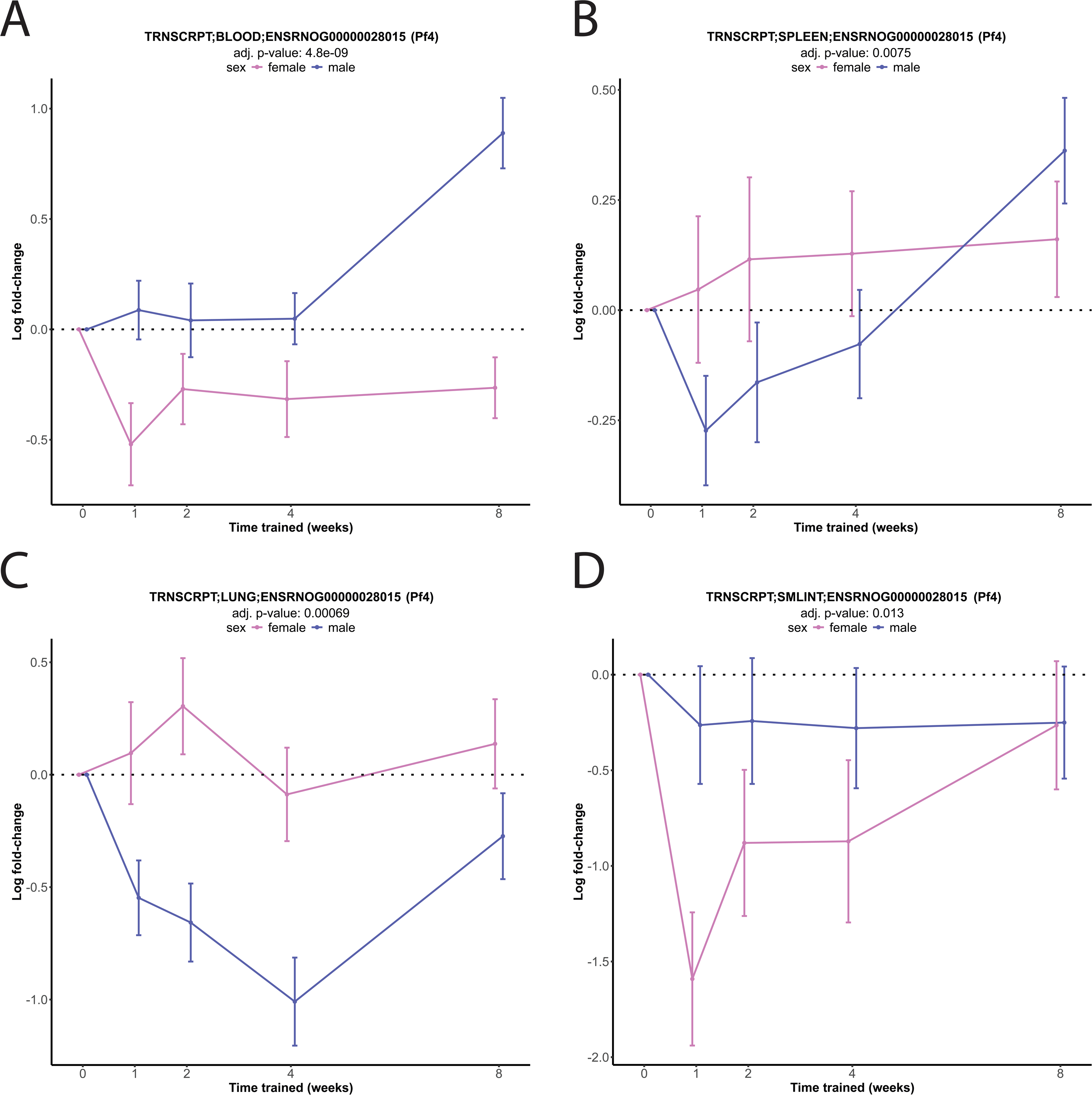
Analysis of platelet factor 4 (*Pf4*) gene expression of 6-month treadmill-exercised rats across all 8 weeks of training in blood cells, spleen, lungs and small intestine (SMLINT) (**A-D**).

#### Gpld1

Gpld1 is an enzyme that plays a role in modulating cell surface dynamics by cleaving glycosylphosphatidylinositols, which affects signaling pathways and cell communication. Acute exercise has been shown to increase Gpld1 levels in the circulation of humans and rodents, which in turn are thought to confer neuroprotective effects [4]. In the kidney, exercise decreased GPLD1 levels in males (p = 0.015) at 1, 4 and 8 weeks, while in females (p < 0.0001), exercise decreased GPLD1 levels across all 8 weeks (**Figure 7A**). GPLD1 WAT levels appeared to decrease for both males and females at 8 weeks, but this was not significant (**Figure 7B**). In BAT, *Gpld1* gene expression decreased only in males (p < 0.0001) across all 8 weeks (**Figure 7C**). In the adrenal gland, *Gpld1* gene expression changed in sex-dimorphic manner: increasing in males (p < 0.0001) at 1, 2 and 8 weeks and decreasing in females (p < 0.0001) across all 8 weeks (**Figure 7D**). Despite GPLD1 being thought of as a liver-originating brain-targeting exerkine [4], exercise did not alter its levels or gene expression in the liver and brain (**Figure S33A-F**). GPLD1 levels were unchanged in gastrocnemius, lungs nor its phosphorylation in the heart (**Figure S33G-I**). *Gpld1* gene expression was unchanged in skeletal muscle (gastrocnemius, vastus lateralis), heart, kidney, colon, spleen, testes, ovaries, lungs, small intestine, blood cells, WAT and vena cava (**Figure S33J-V**).

**Figure 7.**
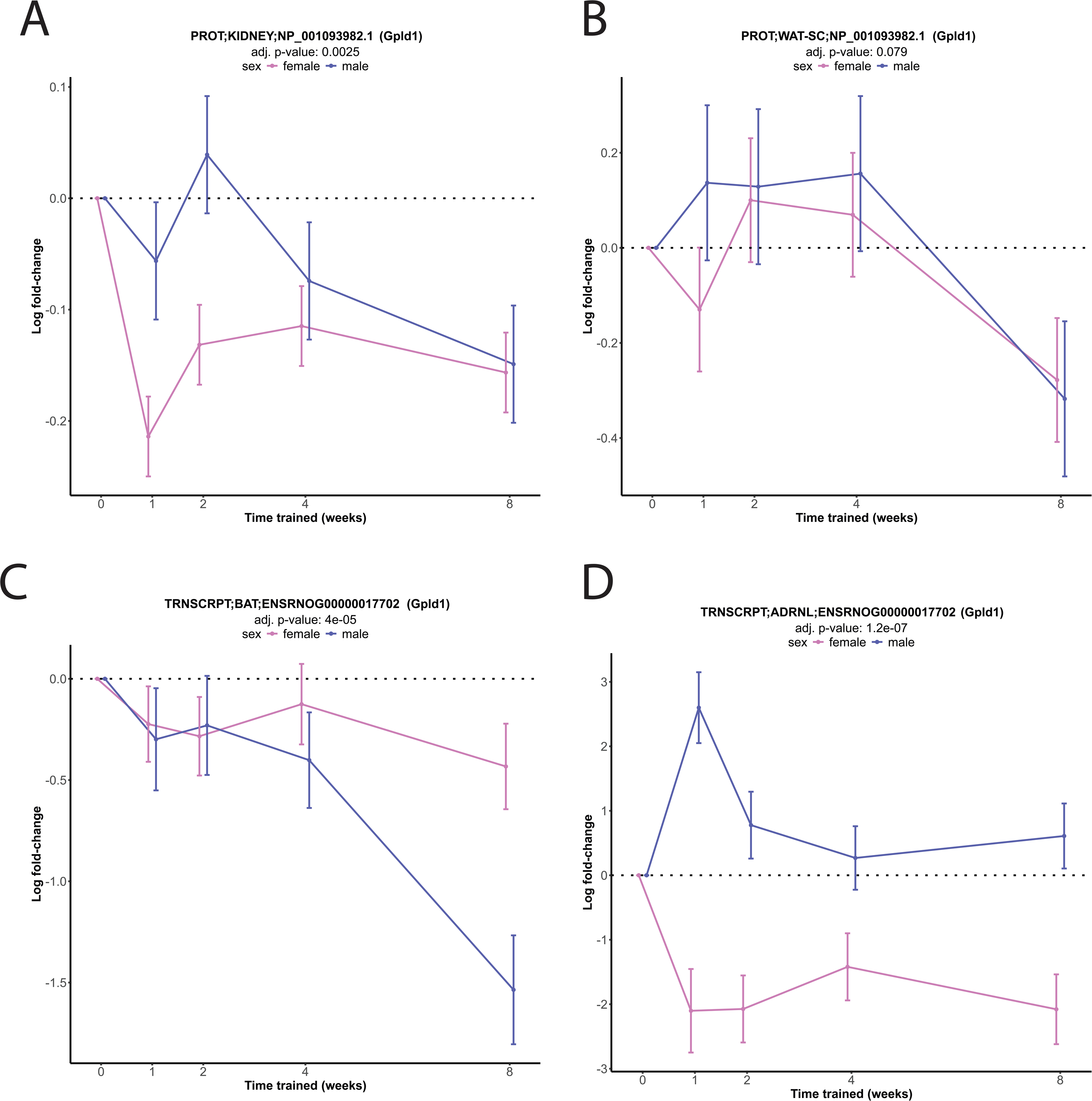
Analysis of glycosylphosphatidylinositol specific phospholipase D1 (GPLD1) levels and gene expression of 6-month treadmill-exercised rats across all 8 weeks of training. GPLD1 levels in the cortex and white adipose tissue (WAT) (**A-B**). *Gpld1* gene expression in brown adipose tissue (BAT) and adrenal gland (ADRNL) (**C-D**).

#### Clusterin

Clusterin, also known as apolipoprotein J, is a glycoprotein implicated in various physiological processes including lipid transport, tissue remodeling, and cell death regulation. It is secreted in response to tissue injury and stress. Recently, its circulation levels have been shown to increase in male rodents with free-access to a wheel and confer neuroprotective and disease-modifying properties against cognitive decline [5].

Even though Clusterin is largely produced by liver hepatocytes, exercise did not alter its levels (including phosphorylation) and gene expression in the liver (**Figure S34A-C)**. Clusterin levels were unchanged in the brain (cortex), skeletal muscle (gastrocnemius), heart, kidney, lungs and WAT (**Figure S34D-I**). Exercise increased *Clu* gene expression in blood cells only in males (p < 0.0001) across all 8 weeks (**Figure 8A**), while it decreased *Clu* gene expression in the vastus lateralis only in females (p < 0.0001) across the same period (**Figure 8B**). In the BAT, *Clu* gene expression decreased only in males (p < 0.0001) at 1 and 2 weeks before suddenly increasing at 8 weeks (**Figure 8C**). In the adrenal gland, *Clu* gene expression increased in males (p = 0.0063) at 1 and 8 weeks and decreased in females (p = 0.01) for the first four weeks (**Figure 8D**). In the colon, *Clu* gene expression increased only in females (p = 0.0003) for the first four weeks (**Figure 8E**). Clusterin phosphorylation levels were unchanged in skeletal muscle (gastrocnemius), the heart, kidney, lungs and WAT (**Figure S34J-O**). *Clu* gene expression was unchanged in the brain (cortex, hippocampus, hypothalamus), skeletal muscle (gastrocnemius, vastus lateralis), heart, kidney, spleen, testes, ovaries, small intestine, BAT, WAT and vena cava (**Figure S34P-ZB**).

**Figure 8.**
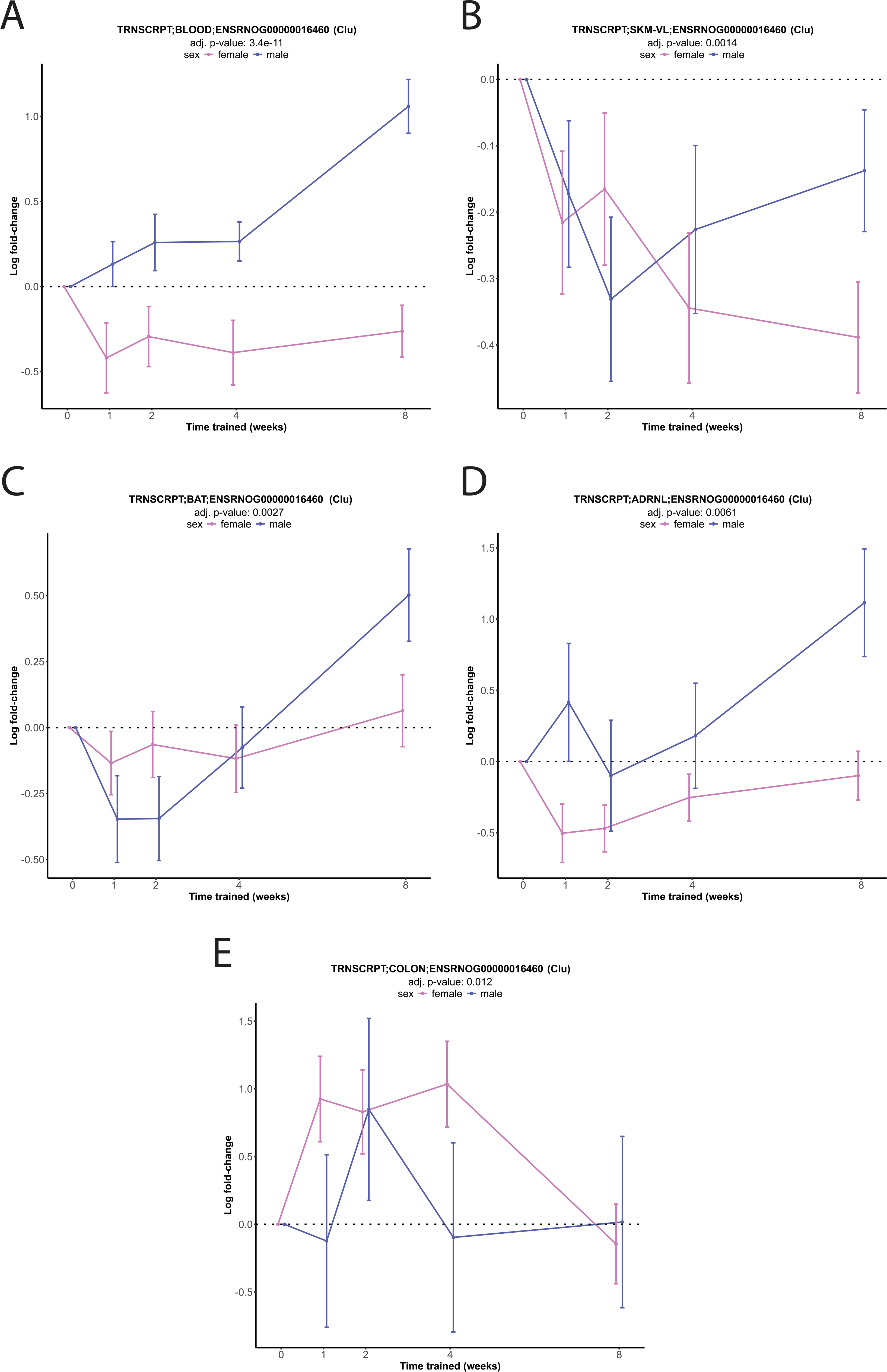
Analysis of Clusterin (*Clu*) gene expression of 6-month treadmill-exercised rats across all 8 weeks of training in blood cells, vastus lateralis (SKM-VL), brown adipose tissue (BAT), adrenal gland and colon (**A-E**).

## Discussion

In this study, we analyzed the levels and transcriptional expression of 26 known exerkines (apelin, adiponectin, METRNL, FGF21, GDF15, FNDC5/irisin, cathepsin B, IL-6, IL-7, IL-10, IL-13, IL-15, musclin, myostatin, LIF, SPARC, SDC4, TGF1β, TGFβ2, angiopoietin1, fractalkine, oxytocin, BDNF, PF4, Gpld1 and Clusterin) and 2 speculative exerkines (PSAP and PSAPL1) across 2 (blood cells and plasma) biofluids and 18 solid tissues (cortex, hippocampus, hypothalamus, gastrocnemius, vastus lateralis. heart, kidney, adrenal gland, colon, spleen, testes, ovaries, lungs, small intestine, liver, BAT, WAT and vena cava) to identify the most exercise-responsive tissues and exerkines. Our analysis revealed distinct exercise-induced patterns of tissue and exerkine responsiveness with time-dependent, intensity-related and sex-dimorphic dynamics.

BAT was the most responsive tissue followed by the adrenal gland, WAT and lungs (**Figure 1A- C**). This is not surprising as exercise is well-documented to alter the molecular signatures of WAT, induce its conversion to BAT (i.e., browning), stimulate thermogenesis, and reduce adipose tissue mass, thereby explaining the observed alterations [131]. However, it was unexpected that popular exerkines known to be upregulated in adipose tissue in response to exercise such as apelin (**Figure S4O-P**) and adiponectin (**Figure S5F-J**) were unchanged. In particular, FNDC5/irisin levels, a very popular WAT-originating exerkine [46], were unchanged in both WAT (**Figure 2B**) and BAT (**Figure S11H**), while its BAT gene expression remained unchanged for 4 weeks and substantially decreased at 8 weeks (**Figure 2F**). Disappointingly, its WAT gene expression also decreased in males across all 8 weeks (**Figure 2G**). It is unclear why this is the case, although there are significant debates surrounding FNDC5/irisin as an exercise-responsive molecule [45].

Surprisingly, despite being frequently studied in the context of exercise, both the brain and plasma showed few exerkine alterations (**Figure 1A**). Specifically, the brain (cortex or hippocampus) only had four exerkine alterations: IL-10 (**Figure S17B**), SPARC (**Figure S23A- B**), fractalkine/CX3CL1 (**Figure 5A-D**), and BDNF (**Figure S30C**). Additionally, all of the popular brain-targeting exerkines including FNDC5/irisin (**Figure S11B-C, L-N**), CTSB (**Figure S12I, Figure S13A-C**), IL-6 (**Figure S15B-C**), Gpld1 (**Figure S33C-F**), Clusterin (**Figure S34D, P-R**) were unchanged in the cortex, hippocampus and/or hypothalamus, while PF4 levels and gene expression were not assessed in the brain. The speculative exerkine PSAP has also been postulated to be brain-targeting [11], however, its levels and gene expression were not altered in the brain (**Figure S14D-H**). Although many studies suggest exercise triggers exerkine release into the circulation [1], only ADIPOQ and BDNF showed changes in plasma: ADIPOQ decreased (**Figure S5A**) and BDNF increased (**Figure S30A-B**), both exclusively in males.

Notably, while prevailing hypotheses suggest skeletal muscle acts as an endocrine organ highly responsive to exercise [60], our findings contrast this view. Specifically, the gastrocnemius had zero exerkine changes, while the vastus lateralis had four exerkine (APLN, **Figure S3A**; PSAP, **Figure 4C**; SPARC, **Figure S23D**; Clusterin, **Figure 8D**) alterations, despite previous studies indicating broader exercise-induced changes in skeletal muscle of humans and rodents [1]. These findings align with a recent organism-wide, 21 cell-type specific plasma secretome analysis of male rodents after one week of treadmill exercise, where the exercise-responsiveness score for skeletal muscle (indicated by molecules released from muscle creatine kinase expressing cells) was relatively low, ranking 12th out of 21 [132]. This also suggests that the vastus lateralis (or generally quadriceps) is more responsive to exercise than the gastrocnemius. Lastly, contrary to recent studies from high-profile journals reporting that the liver releases multiple molecules into the circulation in response to exercise [4; 5; 132], in this study the liver had only 1 exerkine alteration (FNDC5/irisin; **Figure 2E**).

Fractalkine was the most altered exerkine followed by PSAP, CTSB, FNDC5 and SPARC. Fractalkine being the most upregulated exerkine is in agreement with a previous report showing that it had the most elevation in comparison to 11 other myokines in response to low-intensity acute exercise in male humans [133]. Moreover, although PSAP has not previously been identified as an exercise-responsive molecule, the fact that it was altered across seven tissues suggests that future studies should examine its levels and expression in various tissues. This is particularly pertinent given its documented strong neurotrophic effects in the brain [52].

Finally, a key reason for the observed lack of expected exerkine changes in certain tissues may be attributed to biological noise. Tissues are inherently complex and contain various sources of noise such as cellular heterogeneity and varying levels of metabolic activity. Molecules within tissues are also prone to post-translational modifications and enzymatic degradation that can alter protein structures, as well as RNA stability issues such as degradation by RNases and alterations due to ambient RNA-binding proteins.

To circumvent these challenges, one effective method is to measure exerkines in extracellular vesicles (EVs), which are small, bubble-like structures comprising mainly exosomes, ectosomes, and apoptotic bodies, devoid of replication capabilities [134; 135]. EVs are highly responsive to exercise [136–138] and have been extensively utilized in biomarker [139–144] and therapeutic [144–146] studies. Notably, they provide three significant advantages over direct measurement of proteins or RNA in tissues: first, their phospholipid membrane encapsulates and protects their cargo from external modifications or degradation; second, they carry cell-state-specific messages indicating conditions such as stress and disease; and third, their lipophilic nature facilitates easy traversal across cellular membranes. Future studies should explore whether tissue- or exerkine-specific responses differ between bulk tissue quantification vs. their EVs, particularly focusing on time-dependent, intensity-related, and sex-specific variations. Lastly, future studies should explore whether these effects are age-dependent by examining data from 18-month-old rats, and separately determine if these findings are applicable to human subjects once the data is released from the MoTrPaC consortium.

## Additional information

### Data availability

Data used in the preparation of this article were obtained from the Molecular Transducers of Physical Activity Consortium (MoTrPAC) MotrpacRatTraining6moData R package [version 2.0.0].

#### Author contributions

HBT conceptualized and wrote the manuscript.

#### Funding

None.

## Acknowledgements

None.

## REFERENCES

[1] Chow, L.S., Gerszten, R.E., Taylor, J.M., Pedersen, B.K., van Praag, H., Trappe, S., et al., 2022. Exerkines in health, resilience and disease. Nat Rev Endocrinol 18(5):273–289.

[2] Magliulo, L., Bondi, D., Pini, N., Marramiero, L., Di Filippo, E.S., 2022. The wonder exerkines-novel insights: a critical state-of-the-art review. Mol Cell Biochem 477(1):105–113.

[3] Taha, H.B., 2024. Lactate: a potential brain-targeting exerkine. J Physiol.

[4] Horowitz, A.M., Fan, X., Bieri, G., Smith, L.K., Sanchez-Diaz, C.I., Schroer, A.B., et al., 2020. Blood factors transfer beneficial effects of exercise on neurogenesis and cognition to the aged brain. Science 369(6500):167–173.

[5] De Miguel, Z., Khoury, N., Betley, M.J., Lehallier, B., Willoughby, D., Olsson, N., et al., 2021. Exercise plasma boosts memory and dampens brain inflammation via clusterin. Nature 600(7889):494–499.

[6] MoTr, P.A.C.S.G., Lead, A., MoTr, P.A.C.S.G., 2024. Temporal dynamics of the multi-omic response to endurance exercise training. Nature 629(8010):174–183.

[7] Severinsen, M.C.K., Pedersen, B.K., 2020. Muscle-Organ Crosstalk: The Emerging Roles of Myokines. Endocr Rev 41(4):594–609.

[8] Schroer, A.B., Ventura, P.B., Sucharov, J., Misra, R., Chui, M.K.K., Bieri, G., et al., 2023. Platelet factors attenuate inflammation and rescue cognition in ageing. Nature 620(7976):1071–1079.

[9] Leiter, O., Brici, D., Fletcher, S.J., Yong, X.L.H., Widagdo, J., Matigian, N., et al., 2023. Platelet-derived exerkine CXCL4/platelet factor 4 rejuvenates hippocampal neurogenesis and restores cognitive function in aged mice. Nat Commun 14(1):4375.

[10] Mittenbuhler, M.J., Jedrychowski, M.P., Van Vranken, J.G., Sprenger, H.G., Wilensky, S., Dumesic, P.A., et al., 2023. Isolation of extracellular fluids reveals novel secreted bioactive proteins from muscle and fat tissues. Cell Metab 35(3):535–549 e537.

[11] Taha, H.B., Birnbaum, A., Matthews, I., Aceituno, K., Leon, J., Thorwald, M., et al., 2024. Activation of the muscle-to-brain axis ameliorates neurocognitive deficits in an Alzheimer’s disease mouse model via enhancing neurotrophic and synaptic signaling. Geroscience.

[12] Payton, M.E., Greenstone, M.H., Schenker, N., 2003. Overlapping confidence intervals or standard error intervals: what do they mean in terms of statistical significance? J Insect Sci 3:34.

[13] Tatemoto, K., Hosoya, M., Habata, Y., Fujii, R., Kakegawa, T., Zou, M.X., et al., 1998. Isolation and characterization of a novel endogenous peptide ligand for the human APJ receptor. Biochem Biophys Res Commun 251(2):471–476.

[14] Besse-Patin, A., Montastier, E., Vinel, C., Castan-Laurell, I., Louche, K., Dray, C., et al., 2014. Effect of endurance training on skeletal muscle myokine expression in obese men: identification of apelin as a novel myokine. Int J Obes (Lond) 38(5):707–713.

[15] Kwak, S.E., Cho, S.C., Bae, J.H., Lee, J., Shin, H.E., Di Zhang, D., et al., 2019. Effects of exercise-induced apelin on muscle function and cognitive function in aged mice. Exp Gerontol 127:110710.

[16] Vinel, C., Lukjanenko, L., Batut, A., Deleruyelle, S., Pradere, J.P., Le Gonidec, S., et al., 2018. The exerkine apelin reverses age-associated sarcopenia. Nat Med 24(9):1360–1371.

[17] Yan, J., Wang, A., Cao, J., Chen, L., 2020. Apelin/APJ system: an emerging therapeutic target for respiratory diseases. Cell Mol Life Sci 77(15):2919–2930.

[18] Scherer, P.E., Williams, S., Fogliano, M., Baldini, G., Lodish, H.F., 1995. A novel serum protein similar to C1q, produced exclusively in adipocytes. J Biol Chem 270(45):26746–26749.

[19] Fu, Y., Luo, N., Klein, R.L., Garvey, W.T., 2005. Adiponectin promotes adipocyte differentiation, insulin sensitivity, and lipid accumulation. J Lipid Res 46(7):1369–1379.

[20] Yau, S.Y., Li, A., Hoo, R.L., Ching, Y.P., Christie, B.R., Lee, T.M., et al., 2014. Physical exercise-induced hippocampal neurogenesis and antidepressant effects are mediated by the adipocyte hormone adiponectin. Proc Natl Acad Sci U S A 111(44):15810–15815.

[21] Wang, X., You, T., Murphy, K., Lyles, M.F., Nicklas, B.J., 2015. Addition of Exercise Increases Plasma Adiponectin and Release from Adipose Tissue. Med Sci Sports Exerc 47(11):2450–2455.

[22] Schon, M., Kovanicova, Z., Kosutzka, Z., Nemec, M., Tomkova, M., Jackova, L., et al., 2019. Effects of running on adiponectin, insulin and cytokines in cerebrospinal fluid in healthy young individuals. Sci Rep 9(1):1959.

[23] Becic, T., Studenik, C., Hoffmann, G., 2018. Exercise Increases Adiponectin and Reduces Leptin Levels in Prediabetic and Diabetic Individuals: Systematic Review and Meta-Analysis of Randomized Controlled Trials. Med Sci (Basel) 6(4).

[24] Chaolu, H., Asakawa, A., Ushikai, M., Li, Y.X., Cheng, K.C., Li, J.B., et al., 2011. Effect of exercise and high-fat diet on plasma adiponectin and nesfatin levels in mice. Exp Ther Med 2(2):369–373.

[25] Garekani, E.T., Mohebbi, H., Kraemer, R.R., Fathi, R., 2011. Exercise training intensity/volume affects plasma and tissue adiponectin concentrations in the male rat. Peptides 32(5):1008–1012.

[26] Jimenez-Maldonado, A., Virgen-Ortiz, A., Lemus, M., Castro-Rodriguez, E., Cerna-Cortes, J., Muniz, J., et al., 2019. Effects of Moderate- and High-Intensity Chronic Exercise on the Adiponectin Levels in Slow-Twitch and Fast-Twitch Muscles in Rats. Medicina (Kaunas) 55(6).

[27] Martinez-Huenchullan, S.F., Tam, C.S., Ban, L.A., Ehrenfeld-Slater, P., McLennan, S.V., Twigg, S.M., 2020. Skeletal muscle adiponectin induction in obesity and exercise. Metabolism 102:154008.

[28] Zeng, Q., Isobe, K., Fu, L., Ohkoshi, N., Ohmori, H., Takekoshi, K., et al., 2007. Effects of exercise on adiponectin and adiponectin receptor levels in rats. Life Sci 80(5):454–459.

[29] Rizzo, M.R., Fasano, R., Paolisso, G., 2020. Adiponectin and Cognitive Decline. Int J Mol Sci 21(6).

[30] Li, Z., Gao, Z., Sun, T., Zhang, S., Yang, S., Zheng, M., et al., 2023. Meteorin-like/Metrnl, a novel secreted protein implicated in inflammation, immunology, and metabolism: A comprehensive review of preclinical and clinical studies. Front Immunol 14:1098570.

[31] Rao, R.R., Long, J.Z., White, J.P., Svensson, K.J., Lou, J., Lokurkar, I., et al., 2014. Meteorin-like is a hormone that regulates immune-adipose interactions to increase beige fat thermogenesis. Cell 157(6):1279–1291.

[32] Geng, L., Lam, K.S.L., Xu, A., 2020. The therapeutic potential of FGF21 in metabolic diseases: from bench to clinic. Nat Rev Endocrinol 16(11):654–667.

[33] Lin, Z., Tian, H., Lam, K.S., Lin, S., Hoo, R.C., Konishi, M., et al., 2013. Adiponectin mediates the metabolic effects of FGF21 on glucose homeostasis and insulin sensitivity in mice. Cell Metab 17(5):779–789.

[34] Kim, K.H., Kim, S.H., Min, Y.K., Yang, H.M., Lee, J.B., Lee, M.S., 2013. Acute exercise induces FGF21 expression in mice and in healthy humans. PLoS One 8(5):e63517.

[35] Jena, J., Garcia-Pena, L.M., Weatherford, E.T., Marti, A., Bjorkman, S.H., Kato, K., et al., 2023. GDF15 is required for cold-induced thermogenesis and contributes to improved systemic metabolic health following loss of OPA1 in brown adipocytes. Elife 12.

[36] Campderros, L., Moure, R., Cairo, M., Gavalda-Navarro, A., Quesada-Lopez, T., Cereijo, R., et al., 2019. Brown Adipocytes Secrete GDF15 in Response to Thermogenic Activation. Obesity (Silver Spring) 27(10):1606–1616.

[37] Kleinert, M., Clemmensen, C., Sjoberg, K.A., Carl, C.S., Jeppesen, J.F., Wojtaszewski, J.F.P., et al., 2018. Exercise increases circulating GDF15 in humans. Mol Metab 9:187–191.

[38] Klein, A.B., Nicolaisen, T.S., Ortenblad, N., Gejl, K.D., Jensen, R., Fritzen, A.M., et al., 2021. Pharmacological but not physiological GDF15 suppresses feeding and the motivation to exercise. Nat Commun 12(1):1041.

[39] Wrann, C.D., 2015. FNDC5/irisin - their role in the nervous system and as a mediator for beneficial effects of exercise on the brain. Brain Plast 1(1):55–61.

[40] Bostrom, P., Wu, J., Jedrychowski, M.P., Korde, A., Ye, L., Lo, J.C., et al., 2012. A PGC1- alpha-dependent myokine that drives brown-fat-like development of white fat and thermogenesis. Nature 481(7382):463–468.

[41] Hecksteden, A., Wegmann, M., Steffen, A., Kraushaar, J., Morsch, A., Ruppenthal, S., et al., 2013. Irisin and exercise training in humans - results from a randomized controlled training trial. BMC Med 11:235.

[42] Chen, N., Li, Q., Liu, J., Jia, S., 2016. Irisin, an exercise-induced myokine as a metabolic regulator: an updated narrative review. Diabetes Metab Res Rev 32(1):51–59.

[43] Qiu, S., Cai, X., Sun, Z., Schumann, U., Zugel, M., Steinacker, J.M., 2015. Chronic Exercise Training and Circulating Irisin in Adults: A Meta-Analysis. Sports Med 45(11):1577–1588.

[44] Timmons, J.A., Baar, K., Davidsen, P.K., Atherton, P.J., 2012. Is irisin a human exercise gene? Nature 488(7413):E9–10; discussion E10-11.

[45] Albrecht, E., Norheim, F., Thiede, B., Holen, T., Ohashi, T., Schering, L., et al., 2015. Irisin - a myth rather than an exercise-inducible myokine. Sci Rep 5:8889.

[46] Arhire, L.I., Mihalache, L., Covasa, M., 2019. Irisin: A Hope in Understanding and Managing Obesity and Metabolic Syndrome. Front Endocrinol (Lausanne) 10:524.

[47] Islam, M.R., Valaris, S., Young, M.F., Haley, E.B., Luo, R., Bond, S.F., et al., 2021. Exercise hormone irisin is a critical regulator of cognitive function. Nat Metab 3(8):1058–1070.

[48] Nicolini, C., Michalski, B., Toepp, S.L., Turco, C.V., D’Hoine, T., Harasym, D., et al., 2020. A Single Bout of High-intensity Interval Exercise Increases Corticospinal Excitability, Brain-derived Neurotrophic Factor, and Uncarboxylated Osteolcalcin in Sedentary, Healthy Males. Neuroscience 437:242–255.

[49] Gaitan, J.M., Moon, H.Y., Stremlau, M., Dubal, D.B., Cook, D.B., Okonkwo, O.C., et al., 2021. Effects of Aerobic Exercise Training on Systemic Biomarkers and Cognition in Late Middle-Aged Adults at Risk for Alzheimer’s Disease. Front Endocrinol (Lausanne) 12:660181.

[50] Moon, H.Y., Becke, A., Berron, D., Becker, B., Sah, N., Benoni, G., et al., 2016. Running-Induced Systemic Cathepsin B Secretion Is Associated with Memory Function. Cell Metab 24(2):332–340.

[51] O’Brien, J.S., Kretz, K.A., Dewji, N., Wenger, D.A., Esch, F., Fluharty, A.L., 1988. Coding of two sphingolipid activator proteins (SAP-1 and SAP-2) by same genetic locus. Science 241(4869):1098–1101.

[52] Meyer, R.C., Giddens, M.M., Coleman, B.M., Hall, R.A., 2014. The protective role of prosaposin and its receptors in the nervous system. Brain Res 1585:1–12.

[53] Mi, M.Y., Barber, J.L., Rao, P., Farrell, L.A., Sarzynski, M.A., Bouchard, C., et al., 2023. Plasma Proteomic Kinetics in Response to Acute Exercise. Mol Cell Proteomics 22(8):100601.

[54] Kistner, T.M., Pedersen, B.K., Lieberman, D.E., 2022. Interleukin 6 as an energy allocator in muscle tissue. Nat Metab 4(2):170–179.

[55] Tanaka, T., Narazaki, M., Kishimoto, T., 2014. IL-6 in inflammation, immunity, and disease. Cold Spring Harb Perspect Biol 6(10):a016295.

[56] Steensberg, A., van Hall, G., Osada, T., Sacchetti, M., Saltin, B., Klarlund Pedersen, B., 2000. Production of interleukin-6 in contracting human skeletal muscles can account for the exercise-induced increase in plasma interleukin-6. J Physiol 529 Pt 1(Pt 1):237–242.

[57] Docherty, S., Harley, R., McAuley, J.J., Crowe, L.A.N., Pedret, C., Kirwan, P.D., et al., 2022. The effect of exercise on cytokines: implications for musculoskeletal health: a narrative review. BMC Sports Sci Med Rehabil 14(1):5.

[58] Nash, D., Hughes, M.G., Butcher, L., Aicheler, R., Smith, P., Cullen, T., et al., 2023. IL-6 signaling in acute exercise and chronic training: Potential consequences for health and athletic performance. Scand J Med Sci Sports 33(1):4–19.

[59] Haugen, F., Norheim, F., Lian, H., Wensaas, A.J., Dueland, S., Berg, O., et al., 2010. IL-7 is expressed and secreted by human skeletal muscle cells. Am J Physiol Cell Physiol 298(4):C807–816.

[60] Pedersen, B.K., Febbraio, M.A., 2012. Muscles, exercise and obesity: skeletal muscle as a secretory organ. Nat Rev Endocrinol 8(8):457–465.

[61] Cabral-Santos, C., de Lima Junior, E.A., Fernandes, I., Pinto, R.Z., Rosa-Neto, J.C., Bishop, N.C., et al., 2019. Interleukin-10 responses from acute exercise in healthy subjects: A systematic review. J Cell Physiol 234(7):9956–9965.

[62] Nunes, R.B., Tonetto, M., Machado, N., Chazan, M., Heck, T.G., Veiga, A.B., et al., 2008. Physical exercise improves plasmatic levels of IL-10, left ventricular end-diastolic pressure, and muscle lipid peroxidation in chronic heart failure rats. J Appl Physiol (1985) 104(6):1641–1647.

[63] Steensberg, A., Fischer, C.P., Keller, C., Moller, K., Pedersen, B.K., 2003. IL-6 enhances plasma IL-1ra, IL-10, and cortisol in humans. Am J Physiol Endocrinol Metab 285(2):E433–437.

[64] Calegari, L., Nunes, R.B., Mozzaquattro, B.B., Rossato, D.D., Dal Lago, P., 2018. Exercise training improves the IL-10/TNF-alpha cytokine balance in the gastrocnemius of rats with heart failure. Braz J Phys Ther 22(2):154–160.

[65] Wynn, T.A., 2003. IL-13 effector functions. Annu Rev Immunol 21:425–456.

[66] Knudsen, N.H., Stanya, K.J., Hyde, A.L., Chalom, M.M., Alexander, R.K., Liou, Y.H., et al., 2020. Interleukin-13 drives metabolic conditioning of muscle to endurance exercise. Science 368(6490).

[67] Ostrowski, K., Hermann, C., Bangash, A., Schjerling, P., Nielsen, J.N., Pedersen, B.K., 1998. A trauma-like elevation of plasma cytokines in humans in response to treadmill running. J Physiol 513 (Pt 3)(Pt 3):889–894.

[68] Nieman, D.C., Davis, J.M., Henson, D.A., Walberg-Rankin, J., Shute, M., Dumke, C.L., et al., 2003. Carbohydrate ingestion influences skeletal muscle cytokine mRNA and plasma cytokine levels after a 3-h run. J Appl Physiol (1985) 94(5):1917–1925.

[69] Nieman, D.C., Davis, J.M., Brown, V.A., Henson, D.A., Dumke, C.L., Utter, A.C., et al., 2004. Influence of carbohydrate ingestion on immune changes after 2 h of intensive resistance training. J Appl Physiol (1985) 96(4):1292–1298.

[70] Bugera, E.M., Duhamel, T.A., Peeler, J.D., Cornish, S.M., 2018. The systemic myokine response of decorin, interleukin-6 (IL-6) and interleukin-15 (IL-15) to an acute bout of blood flow restricted exercise. Eur J Appl Physiol 118(12):2679–2686.

[71] Nishida, Y., Tanaka, K., Hara, M., Hirao, N., Tanaka, H., Tobina, T., et al., 2015. Effects of home-based bench step exercise on inflammatory cytokines and lipid profiles in elderly Japanese females: A randomized controlled trial. Arch Gerontol Geriatr 61(3):443–451.

[72] Roh, H.T., Cho, S.Y., So, W.Y., 2020. Effects of Regular Taekwondo Intervention on Oxidative Stress Biomarkers and Myokines in Overweight and Obese Adolescents. Int J Environ Res Public Health 17(7).

[73] Perez-Lopez, A., McKendry, J., Martin-Rincon, M., Morales-Alamo, D., Perez-Kohler, B., Valades, D., et al., 2018. Skeletal muscle IL-15/IL-15Ralpha and myofibrillar protein synthesis after resistance exercise. Scand J Med Sci Sports 28(1):116–125.

[74] Oliver, J.M., Jenke, S.C., Mata, J.D., Kreutzer, A., Jones, M.T., 2016. Acute Effect of Cluster and Traditional Set Configurations on Myokines Associated with Hypertrophy. Int J Sports Med 37(13):1019–1024.

[75] Park, K.M., Park, S.C., Kang, S., 2019. Effects of resistance exercise on adipokine factors and body composition in pre- and postmenopausal women. J Exerc Rehabil 15(5):676–682.

[76] Quinn, L.S., Anderson, B.G., Conner, J.D., Wolden-Hanson, T., Marcell, T.J., 2014. IL-15 is required for postexercise induction of the pro-oxidative mediators PPARdelta and SIRT1 in male mice. Endocrinology 155(1):143–155.

[77] Minuzzi, L.G., da Conceicao, L.R., Munoz, V.R., Vieira, R.F.L., Gaspar, R.C., da Silva, A.S.R., et al., 2021. Effects of short-term physical training on the interleukin-15 signalling pathway and glucose tolerance in aged rats. Cytokine 137:155306.

[78] Alvarez, B., Carbo, N., Lopez-Soriano, J., Drivdahl, R.H., Busquets, S., Lopez-Soriano, F.J., et al., 2002. Effects of interleukin-15 (IL-15) on adipose tissue mass in rodent obesity models: evidence for direct IL-15 action on adipose tissue. Biochim Biophys Acta 1570(1):33–37.

[79] Carbo, N., Lopez-Soriano, J., Costelli, P., Alvarez, B., Busquets, S., Baccino, F.M., et al., 2001. Interleukin-15 mediates reciprocal regulation of adipose and muscle mass: a potential role in body weight control. Biochim Biophys Acta 1526(1):17–24.

[80] Yang, H., Chang, J., Chen, W., Zhao, L., Qu, B., Tang, C., et al., 2013. Treadmill exercise promotes interleukin 15 expression in skeletal muscle and interleukin 15 receptor alpha expression in adipose tissue of high-fat diet rats. Endocrine 43(3):579–585.

[81] Nishizawa, H., Matsuda, M., Yamada, Y., Kawai, K., Suzuki, E., Makishima, M., et al., 2004. Musclin, a novel skeletal muscle-derived secretory factor. J Biol Chem 279(19):19391–19395.

[82] Subbotina, E., Sierra, A., Zhu, Z., Gao, Z., Koganti, S.R., Reyes, S., et al., 2015. Musclin is an activity-stimulated myokine that enhances physical endurance. Proc Natl Acad Sci U S A 112(52):16042–16047.

[83] Nam, J.S., Cho, E.S., Kwon, Y.R., Park, J.S., Kim, Y., 2024. Dynamic Response of Musclin, a Myokine, to Aerobic Exercise and Its Interplay with Natriuretic Peptides and Receptor C. J Clin Endocrinol Metab.

[84] McPherron, A.C., Lawler, A.M., Lee, S.J., 1997. Regulation of skeletal muscle mass in mice by a new TGF-beta superfamily member. Nature 387(6628):83–90.

[85] Shan, T., Liang, X., Bi, P., Kuang, S., 2013. Myostatin knockout drives browning of white adipose tissue through activating the AMPK-PGC1alpha-Fndc5 pathway in muscle. FASEB J 27(5):1981–1989.

[86] Jin, L., Han, S., Lv, X., Li, X., Zhang, Z., Kuang, H., et al., 2023. The muscle-enriched myokine Musclin impairs beige fat thermogenesis and systemic energy homeostasis via Tfr1/PKA signaling in male mice. Nat Commun 14(1):4257.

[87] Zhang, C., McFarlane, C., Lokireddy, S., Masuda, S., Ge, X., Gluckman, P.D., et al., 2012. Inhibition of myostatin protects against diet-induced obesity by enhancing fatty acid oxidation and promoting a brown adipose phenotype in mice. Diabetologia 55(1):183–193.

[88] Jorgensen, M.M., de la Puente, P., 2022. Leukemia Inhibitory Factor: An Important Cytokine in Pathologies and Cancer. Biomolecules 12(2).

[89] Broholm, C., Pedersen, B.K., 2010. Leukaemia inhibitory factor--an exercise-induced myokine. Exerc Immunol Rev 16:77–85.

[90] Broholm, C., Mortensen, O.H., Nielsen, S., Akerstrom, T., Zankari, A., Dahl, B., et al., 2008. Exercise induces expression of leukaemia inhibitory factor in human skeletal muscle. J Physiol 586(8):2195–2201.

[91] Zouein, F.A., Kurdi, M., Booz, G.W., 2013. LIF and the heart: just another brick in the wall? Eur Cytokine Netw 24(1):11–19.

[92] Jia, D., Cai, M., Xi, Y., Du, S., ZhenjunTian, 2018. Interval exercise training increases LIF expression and prevents myocardial infarction-induced skeletal muscle atrophy in rats. Life Sci 193:77–86.

[93] Aoi, W., Naito, Y., Takagi, T., Tanimura, Y., Takanami, Y., Kawai, Y., et al., 2013. A novel myokine, secreted protein acidic and rich in cysteine (SPARC), suppresses colon tumorigenesis via regular exercise. Gut 62(6):882–889.

[94] Ghanemi, A., Yoshioka, M., St-Amand, J., 2021. Measuring Exercise-Induced Secreted Protein Acidic and Rich in Cysteine Expression as a Molecular Tool to Optimize Personalized Medicine. Genes (Basel) 12(11).

[95] Ronning, S.B., Carlson, C.R., Stang, E., Kolset, S.O., Hollungs, K., Pedersen, M.E., 2015. Syndecan-4 Regulates Muscle Differentiation and Is Internalized from the Plasma Membrane during Myogenesis. PLoS One 10(6):e0129288.

[96] Lee, S., Kolset, S.O., Birkeland, K.I., Drevon, C.A., Reine, T.M., 2019. Acute exercise increases syndecan-1 and −4 serum concentrations. Glycoconj J 36(2):113–125.

[97] De Nardo, W., Miotto, P.M., Bayliss, J., Nie, S., Keenan, S.N., Montgomery, M.K., et al., 2022. Proteomic analysis reveals exercise training induced remodelling of hepatokine secretion and uncovers syndecan-4 as a regulator of hepatic lipid metabolism. Mol Metab 60:101491.

[98] Morikawa, M., Derynck, R., Miyazono, K., 2016. TGF-beta and the TGF-beta Family: Context-Dependent Roles in Cell and Tissue Physiology. Cold Spring Harb Perspect Biol 8(5).

[99] Li, Y., Foster, W., Deasy, B.M., Chan, Y., Prisk, V., Tang, Y., et al., 2004. Transforming growth factor-beta1 induces the differentiation of myogenic cells into fibrotic cells in injured skeletal muscle: a key event in muscle fibrogenesis. Am J Pathol 164(3):1007–1019.

[100] Heinemeier, K.M., Bjerrum, S.S., Schjerling, P., Kjaer, M., 2013. Expression of extracellular matrix components and related growth factors in human tendon and muscle after acute exercise. Scand J Med Sci Sports 23(3):e150–161.

[101] Breen, E.C., Johnson, E.C., Wagner, H., Tseng, H.M., Sung, L.A., Wagner, P.D., 1996. Angiogenic growth factor mRNA responses in muscle to a single bout of exercise. J Appl Physiol (1985) 81(1):355–361.

[102] Takahashi, H., Alves, C.R.R., Stanford, K.I., Middelbeek, R.J.W., Nigro, P., Ryan, R.E., et al., 2019. TGF-beta2 is an exercise-induced adipokine that regulates glucose and fatty acid metabolism. Nat Metab 1(2):291–303.

[103] Ding, Y.H., Li, J., Zhou, Y., Rafols, J.A., Clark, J.C., Ding, Y., 2006. Cerebral angiogenesis and expression of angiogenic factors in aging rats after exercise. Curr Neurovasc Res 3(1):15–23.

[104] Cullberg, K.B., Christiansen, T., Paulsen, S.K., Bruun, J.M., Pedersen, S.B., Richelsen, B., 2013. Effect of weight loss and exercise on angiogenic factors in the circulation and in adipose tissue in obese subjects. Obesity (Silver Spring) 21(3):454–460.

[105] Gavin, T.P., Drew, J.L., Kubik, C.J., Pofahl, W.E., Hickner, R.C., 2007. Acute resistance exercise increases skeletal muscle angiogenic growth factor expression. Acta Physiol (Oxf) 191(2):139–146.

[106] Catoire, M., Mensink, M., Kalkhoven, E., Schrauwen, P., Kersten, S., 2014. Identification of human exercise-induced myokines using secretome analysis. Physiol Genomics 46(7):256–267.

[107] Wong, B.W., Wong, D., McManus, B.M., 2002. Characterization of fractalkine (CX3CL1) and CX3CR1 in human coronary arteries with native atherosclerosis, diabetes mellitus, and transplant vascular disease. Cardiovasc Pathol 11(6):332–338.

[108] Kumar, P., Stiernborg, M., Fogdell-Hahn, A., Mansson, K., Furmark, T., Berglind, D., et al., 2022. Physical exercise is associated with a reduction in plasma levels of fractalkine, TGF-beta1, eotaxin-1 and IL-6 in younger adults with mobility disability. PLoS One 17(2):e0263173.

[109] Farrash, W.F., Phillips, B.E., Britton, S.L., Qi, N., Koch, L.G., Wilkinson, D.J., et al., 2021. Myokine Responses to Exercise in a Rat Model of Low/High Adaptive Potential. Front Endocrinol (Lausanne) 12:645881.

[110] Elabd, C., Cousin, W., Upadhyayula, P., Chen, R.Y., Chooljian, M.S., Li, J., et al., 2014. Oxytocin is an age-specific circulating hormone that is necessary for muscle maintenance and regeneration. Nat Commun 5:4082.

[111] Tsukamoto, H., Olesen, N.D., Petersen, L.G., Suga, T., Sorensen, H., Nielsen, H.B., et al., 2024. Circulating Plasma Oxytocin Level Is Elevated by High-Intensity Interval Exercise in Men. Med Sci Sports Exerc 56(5):927–932.

[112] Rassovsky, Y., Harwood, A., Zagoory-Sharon, O., Feldman, R., 2019. Martial arts increase oxytocin production. Sci Rep 9(1):12980.

[113] Yuksel, O., Ates, M., Kizildag, S., Yuce, Z., Koc, B., Kandis, S., et al., 2019. Regular Aerobic Voluntary Exercise Increased Oxytocin in Female Mice: The Cause of Decreased Anxiety and Increased Empathy-Like Behaviors. Balkan Med J 36(5):257–262.

[114] Miranda, M., Morici, J.F., Zanoni, M.B., Bekinschtein, P., 2019. Brain-Derived Neurotrophic Factor: A Key Molecule for Memory in the Healthy and the Pathological Brain. Front Cell Neurosci 13:363.

[115] Reycraft, J.T., Islam, H., Townsend, L.K., Hayward, G.C., Hazell, T.J., Macpherson, R.E.K., 2020. Exercise Intensity and Recovery on Circulating Brain-derived Neurotrophic Factor. Med Sci Sports Exerc 52(5):1210–1217.

[116] Voss, M.W., Soto, C., Yoo, S., Sodoma, M., Vivar, C., van Praag, H., 2019. Exercise and Hippocampal Memory Systems. Trends Cogn Sci 23(4):318–333.

[117] Yau, S.Y., Lau, B.W., Zhang, E.D., Lee, J.C., Li, A., Lee, T.M., et al., 2012. Effects of voluntary running on plasma levels of neurotrophins, hippocampal cell proliferation and learning and memory in stressed rats. Neuroscience 222:289–301.

[118] Goekint, M., Bos, I., Heyman, E., Meeusen, R., Michotte, Y., Sarre, S., 2012. Acute running stimulates hippocampal dopaminergic neurotransmission in rats, but has no influence on brain-derived neurotrophic factor. J Appl Physiol (1985) 112(4):535–541.

[119] Liu, P.Z., Nusslock, R., 2018. Exercise-Mediated Neurogenesis in the Hippocampus via BDNF. Front Neurosci 12:52.

[120] Barha, C.K., Falck, R.S., Davis, J.C., Nagamatsu, L.S., Liu-Ambrose, T., 2017. Sex differences in aerobic exercise efficacy to improve cognition: A systematic review and meta-analysis of studies in older rodents. Front Neuroendocrinol 46:86–105.

[121] Cuppini, R., Sartini, S., Agostini, D., Guescini, M., Ambrogini, P., Betti, M., et al., 2007. Bdnf expression in rat skeletal muscle after acute or repeated exercise. Arch Ital Biol 145(2):99–110.

[122] Renteria, I., Garcia-Suarez, P.C., Fry, A.C., Moncada-Jimenez, J., Machado-Parra, J.P., Antunes, B.M., et al., 2022. The Molecular Effects of BDNF Synthesis on Skeletal Muscle: A Mini-Review. Front Physiol 13:934714.

[123] Mousavi, K., Jasmin, B.J., 2006. BDNF is expressed in skeletal muscle satellite cells and inhibits myogenic differentiation. J Neurosci 26(21):5739–5749.

[124] Delezie, J., Weihrauch, M., Maier, G., Tejero, R., Ham, D.J., Gill, J.F., et al., 2019. BDNF is a mediator of glycolytic fiber-type specification in mouse skeletal muscle. Proc Natl Acad Sci U S A 116(32):16111–16120.

[125] Stanford, K.I., Goodyear, L.J., 2016. Exercise regulation of adipose tissue. Adipocyte 5(2):153–162.

[126] Stanford, K.I., Middelbeek, R.J., Goodyear, L.J., 2015. Exercise Effects on White Adipose Tissue: Beiging and Metabolic Adaptations. Diabetes 64(7):2361–2368.

[127] Dhaliwal, R., 2023. Examining the effects of BDNF and exercise training on adipose tissue browning Faculty of Applied Health Sciences. Brock University.

[128] Villeda, S.A., Plambeck, K.E., Middeldorp, J., Castellano, J.M., Mosher, K.I., Luo, J., et al., 2014. Young blood reverses age-related impairments in cognitive function and synaptic plasticity in mice. Nat Med 20(6):659–663.

[129] Dubal, D.B., Yokoyama, J.S., Zhu, L., Broestl, L., Worden, K., Wang, D., et al., 2014. Life extension factor klotho enhances cognition. Cell Rep 7(4):1065–1076.

[130] Leiter, O., Seidemann, S., Overall, R.W., Ramasz, B., Rund, N., Schallenberg, S., et al., 2019. Exercise-Induced Activated Platelets Increase Adult Hippocampal Precursor Proliferation and Promote Neuronal Differentiation. Stem Cell Reports 12(4):667–679.

[131] Aldiss, P., Betts, J., Sale, C., Pope, M., Budge, H., Symonds, M.E., 2018. Exercise-induced ‘browning’ of adipose tissues. Metabolism 81:63–70.

[132] Wei, W., Riley, N.M., Lyu, X., Shen, X., Guo, J., Raun, S.H., et al., 2023. Organism-wide, cell-type-specific secretome mapping of exercise training in mice. Cell Metab 35(7):1261–1279 e1211.

[133] Hashida, R., Matsuse, H., Kawaguchi, T., Yoshio, S., Bekki, M., Iwanaga, S., et al., 2021. Effects of a low-intensity resistance exercise program on serum miR-630, miR-5703, and Fractalkine/CX3CL1 expressions in subjects with No exercise habits: A preliminary study. Hepatol Res 51(7):823–833.

[134] van Niel, G., D’Angelo, G., Raposo, G., 2018. Shedding light on the cell biology of extracellular vesicles. Nat Rev Mol Cell Biol 19(4):213–228.

[135] Dixson, A.C., Dawson, T.R., Di Vizio, D., Weaver, A.M., 2023. Context-specific regulation of extracellular vesicle biogenesis and cargo selection. Nat Rev Mol Cell Biol 24(7):454–476.

[136] Nederveen, J.P., Warnier, G., Di Carlo, A., Nilsson, M.I., Tarnopolsky, M.A., 2020. Extracellular Vesicles and Exosomes: Insights From Exercise Science. Front Physiol 11:604274.

[137] Vechetti, I.J., Jr., Valentino, T., Mobley, C.B., McCarthy, J.J., 2021. The role of extracellular vesicles in skeletal muscle and systematic adaptation to exercise. J Physiol 599(3):845–861.

[138] Whitham, M., Parker, B.L., Friedrichsen, M., Hingst, J.R., Hjorth, M., Hughes, W.E., et al., 2018. Extracellular Vesicles Provide a Means for Tissue Crosstalk during Exercise. Cell Metab 27(1):237–251 e234.

[139] Taha, H.B., Bogoniewski, A., 2024. Analysis of biomarkers in speculative CNS-enriched extracellular vesicles for parkinsonian disorders: a comprehensive systematic review and diagnostic meta-analysis. J Neurol 271(4):1680–1706.

[140] Taha, H.B., Bogoniewski, A., 2023. Extracellular vesicles from bodily fluids for the accurate diagnosis of Parkinson’s disease and related disorders: A systematic review and diagnostic meta-analysis. J Extracell Biol 2(11):e121.

[141] Dutta, S., Hornung, S., Taha, H.B., Bitan, G., 2023. Biomarkers for parkinsonian disorders in CNS-originating EVs: promise and challenges. Acta Neuropathol 145(5):515–540.

[142] Urabe, F., Kosaka, N., Ito, K., Kimura, T., Egawa, S., Ochiya, T., 2020. Extracellular vesicles as biomarkers and therapeutic targets for cancer. Am J Physiol Cell Physiol 318(1):C29–C39.

[143] Huang-Doran, I., Zhang, C.Y., Vidal-Puig, A., 2017. Extracellular Vesicles: Novel Mediators of Cell Communication In Metabolic Disease. Trends Endocrinol Metab 28(1):3–18.

[144] Khadka, A., Spiers, J.G., Cheng, L., Hill, A.F., 2023. Extracellular vesicles with diagnostic and therapeutic potential for prion diseases. Cell Tissue Res 392(1):247–267.

[145] Pait, M.C., Kaye, S.D., Su, Y., Kumar, A., Singh, S., Gironda, S.C., et al., 2024. Novel method for collecting hippocampal interstitial fluid extracellular vesicles (EV(ISF)) reveals sex-dependent changes in microglial EV proteome in response to Abeta pathology. J Extracell Vesicles 13(1):e12398.

[146] Lener, T., Gimona, M., Aigner, L., Borger, V., Buzas, E., Camussi, G., et al., 2015. Applying extracellular vesicles based therapeutics in clinical trials - an ISEV position paper. J Extracell Vesicles 4:30087.

